# Scalable subclonal reconstruction of cancer cells in DNA sequencing data using a penalized likelihood model

**DOI:** 10.1101/2021.03.31.437383

**Authors:** Yujie Jiang, Matthew D Montierth, Yu Ding, Kaixian Yu, Quang Tran, Aaron Wu, Ruonan Li, Shuangxi Ji, Xiaoqian Liu, Seung Jun Shin, Shaolong Cao, Yuxin Tang, Tom Lesluyes, Marek Kimmel, Jennifer R. Wang, Maxime Tarabichi, Hongtu Zhu, Peter Van Loo, Wenyi Wang

## Abstract

**Background:** Characterizing intratumor heterogeneity and subclonal architecture from bulk DNA sequencing provides important insights into cancer evolution. However, existing subclonal reconstruction methods remain computationally demanding, hindering iterative quality control and reproducible application across large genomic cohorts.

**Methods:** Distinct from existing likelihood-based and Bayesian approaches, we developed CliPP (Clonal structure identification through Pairwise Penalization), utilizing machine learning to add pairwise fusion penalty to the full likelihood, in order to simultaneously estimate mutation cellular prevalence and fuse compatible estimates into clusters without prespecifying the number of clusters. We benchmarked CliPP in three simulation datasets comprising 5,256 tumors for accuracy, and further evaluated its concordance with consensus calls in 2,008 tumors profiled by whole-genome sequencing from the Pan-Cancer Analysis of Whole Genomes (PCAWG) study. We then applied a standardized, quality-controlled workflow to whole-exome sequencing data from The Cancer Genome Atlas (TCGA), yielding a high-confidence cohort of 5,424 tumors across 31 cancer types. Within this cohort, we mapped the clonality of 13,960 high-confidence driver mutations in 4,375 tumors.

**Results:** CliPP showed competitive performance across all datasets, as compared to PhylogicNDT and PyClone-VI, while gaining significant improvement in speed. In PCAWG, CliPP achieved a concordance correlation coefficient (CCC) of 0.93 with the published consensus, leading to a 96.3% agreement in driver-level clonal/subclonal assignments from the PCAWG study (Cohen’s κ = 0.81). It took approximately one hour for CliPP to complete the analysis of the entire PCAWG and TCGA cohorts, gaining noticeable time over existing methods. Applying CliPP to the TCGA cohort, we generate a complete annotation of driver clonality across TCGA samples, and observe broad patterns of driver timing that recapitulate known biology. The CliPP-on-Web app allows non-computational researchers to directly obtain subclonal annotations of their tumor samples.

**Conclusions:** Machine learning assisted CliPP combines competitive reconstruction accuracy, substantial computational efficiency, and standardized quality control to enable reproducible subclonal reconstruction at large scale. Its pan-cancer application characterizes context-dependent patterns of driver clonality, and its software web interface enables cancer evolution inquiries from the cancer research community.

## Background

Mutations in tumor cells accumulate over time, starting from a single cancer-initiating cell and progressing to the diverse mutations present across a mature tumor [1,2]. This process leads to the development of heterogeneous subpopulations of cells, i.e., (sub)clones, with unique genetic profiles from the ancestral cancer cell [3–5] (**Fig. 1a**). Over the past decade, lineage-resolved studies have advanced our understanding of the temporal and spatial dynamics of mutation acquisition across tumor clones [6–11], with many supported by pan-cancer investigations [12–17]. Insights from these studies have created new opportunities for the development of cancer therapies [18,19].

**Figure 1:**
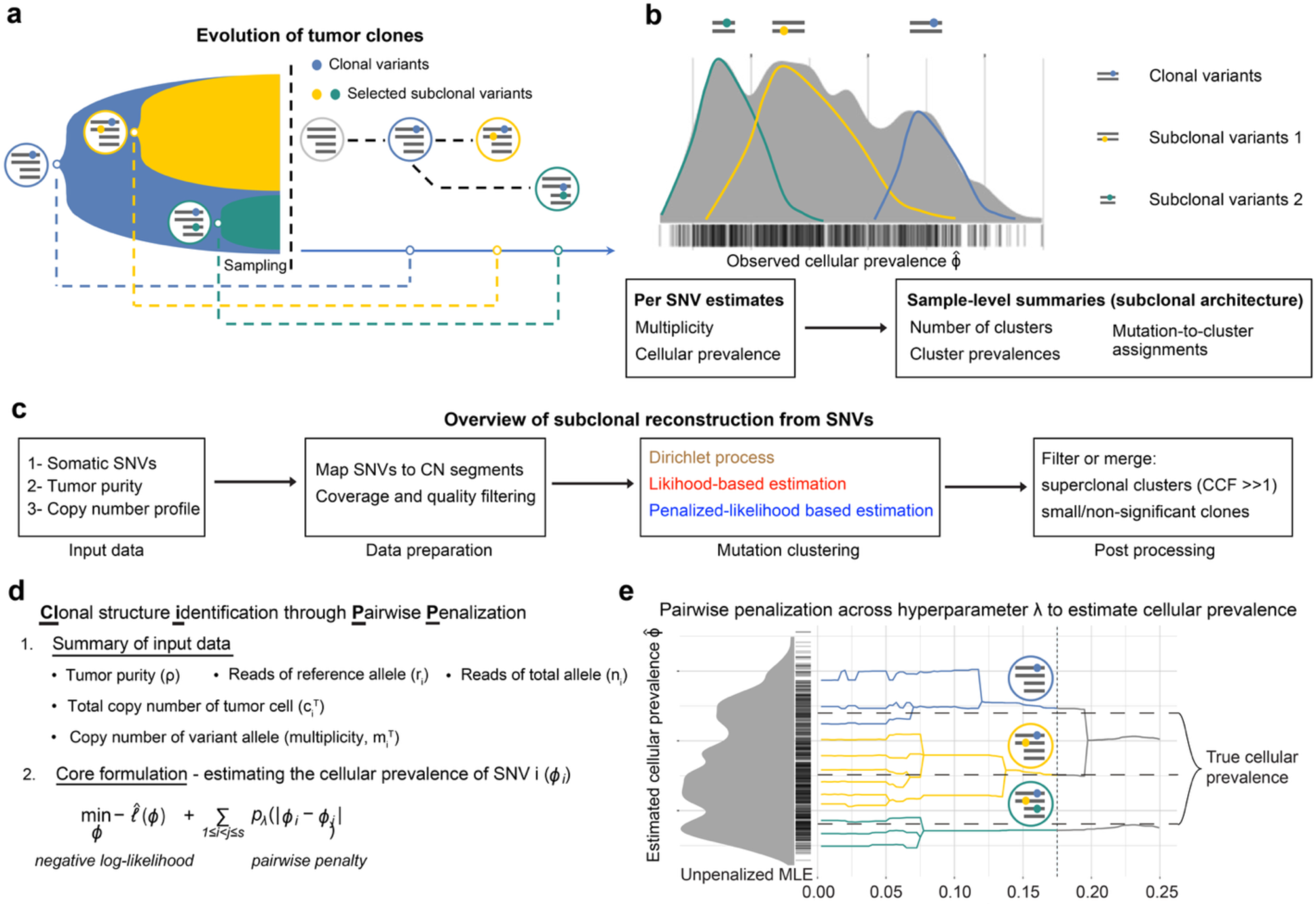
Subclonal reconstruction and CliPP overview. **(a)** A visual representation of tumor evolution with genotype circles indicating representative somatic single nucleotide variants (SNVs). Colored dots represent SNVs. **(b)** Cellular prevalence (CP) distribution of SNVs (each SNV is a bar along the X axis) for an illustrative example, characterized by one clonal cluster (blue) and two subclonal clusters (yellow and green). Multiplicity: SNV-specific copy number, which can be different from allele-specific copy number. **(c)** An overview of SNV-based subclonal reconstruction. Somatic SNVs, SNV-specific VAFs, tumor purity and copy-number profiles are first harmonized through preprocessing and quality control, then followed by mutation clustering using representative classes of statistical models. Existing approaches include Dirichlet-process models (brown) and likelihood-based methods (red), whereas CliPP uses penalized-likelihood estimation (blue). Post-processing merges implausible “superclusters” (e.g., clusters with estimated CCF >> 1) that are often caused by inaccurate copy number multiplicity input, and insignificant clones/subclones. This pipeline yields refined subclonal architectures that are used for downstream phylogenetic/evolutionary analyses, pan-cancer subclonal landscape profiling, driver gene analyses, etc. **(d)** Input data and the core optimization problem for the CliPP model. **(e)** CliPP infers clusters of SNVs and cellular prevalence (CP) values using a penalty function that varies over a penalty parameter, λ, which is shown along the x-axis. The y-axis corresponds to CP estimates (0 to 1) connecting the nearest CP estimates over 100 evenly spaced λ values (0.01 to 0.25). The vertical dotted line indicates the λ corresponding to the identified optimal solution (see **Methods**). Somatic SNV clusters distinguish clonal cancerous cells (blue) from new subclonal cancerous cell populations (yellow and green) that converge for homogeneity pursuit in CPs.

Subclonal reconstruction from bulk tumor sequencing seeks to recover this heterogeneous architecture by clustering somatic single-nucleotide variants (SNVs) according to their cellular prevalence. In practice, variant read counts and variant allele fractions are modeled jointly with sample-specific purity and allele-specific copy-number states so that mutations arising in the same subclone share similar cellular prevalence and can be grouped together (**Fig. 1b**). These clustered mutations then provide a quantitative summary of a tumor’s clonal and subclonal structure and support downstream analyses of tumor phylogeny and evolutionary timing.

Despite valuable insights from cancer-type specific, deeply annotated cohorts [10,20–23], large-scale translational studies of tumor evolutionary dynamics remain challenging due to computational burden, sensitivity to modeling and input data quality, and the frequent need for per-sample manual curation of input/output for subclonal inference [13]. These challenges highlight the need for scalable, efficient, and accurate subclonal reconstruction methods to advance cancer evolution research. Widely used Bayesian mixture-model and Dirichlet-process frameworks (e.g., PyClone [24], PhyloWGS [25], DPClust [26] and extensions) integrate variant allele fractions with purity and copy-number profiles to infer mutation clusters and sometimes phylogenies, but typically require priors or their surrogates, intensive MCMC, careful hyperparameter tuning, and non-trivial post hoc curation (e.g., merging/pruning clusters). These demands, together with inter- and intra-tumor heterogeneity and per-sample manual curation, make analyses computationally expensive, particularly for whole-genome sequencing data, sensitive to implementation details, and challenging to apply reproducibly across cohorts of thousands of tumors or in iterative workflows [4,21]. More recently, PhylogicNDT [27] and PyClone-VI [28] deployed advanced techniques including variational inference to substantially speed up the Bayesian methods, without changing the core estimation process.

Current single-sample subclonal reconstruction methods also show room for improvement in recovering basic characteristics such as the proportion of mutations classified as subclonal (subclonal fraction), ignoring the further clustering structure among subclonal mutations [13]. While multi-sample sequencing provides a solution for improved accuracy, its corresponding multi-sample based mutation clustering is more challenging, while there are tens of thousands of patient samples with only single-sample DNA sequencing that need to be mined [12,30–33]. Therefore, approaching subclonal reconstruction from an orthogonal parameter estimation perspective may offer opportunities to improve reconstruction accuracy, potentially gaining computational speed, and preserving both advantages for future extension to multi-sample sequencing.

To address these limitations, we developed **CliPP** (**Clonal structure identification through Pairwise Penalization)**, a method using a modeling approach orthogonal to previous methods: regularized maximum likelihood, for subclonal reconstruction from bulk DNA sequencing data (**Fig. 1c-e**). In this study, we benchmark the accuracy and speed against existing approaches using extensive simulations and whole-genome sequencing (WGS) data from 2,008 patient tumor samples in the International Cancer Genome Consortium Pan Cancer Analysis of Whole Genomes (ICGC-PCAWG). We then applied a standardized, quality-controlled workflow to whole-exome sequencing (WES) data from TCGA, characterizing subclonal architecture across 5,424 tumors from 31 cancer types and mapping the clonality of 13,960 high-confidence driver mutations in 4,375 tumors. CliPP’s superior computational speed enables a deployment as a user friendly Shiny app, which is freely available for cancer researchers at: https://bioinformatics.mdanderson.org/apps/CliPP [34].

## Methods

### The CliPP Model

In the wake of the genomic revolution, several approaches have been developed to call and quantify cancer cell clones through clustering of variant read counts [35]. Many of the existing methods address subclonal reconstruction with mixture models, as implemented via Dirichlet processes [36], which are computationally costly, especially when the underlying subclonal structure is complex. CliPP presents a distinct statistical framework that addresses this task through penalized regression, in order to reduce the computational burden while maintaining high accuracy. It ensures homogeneity by introducing a pairwise penalty on parameter estimation without the need to pre-order coefficients, an advancement over preceding methods such as fused LASSO [37] and CARDS algorithm [38].

#### Notations

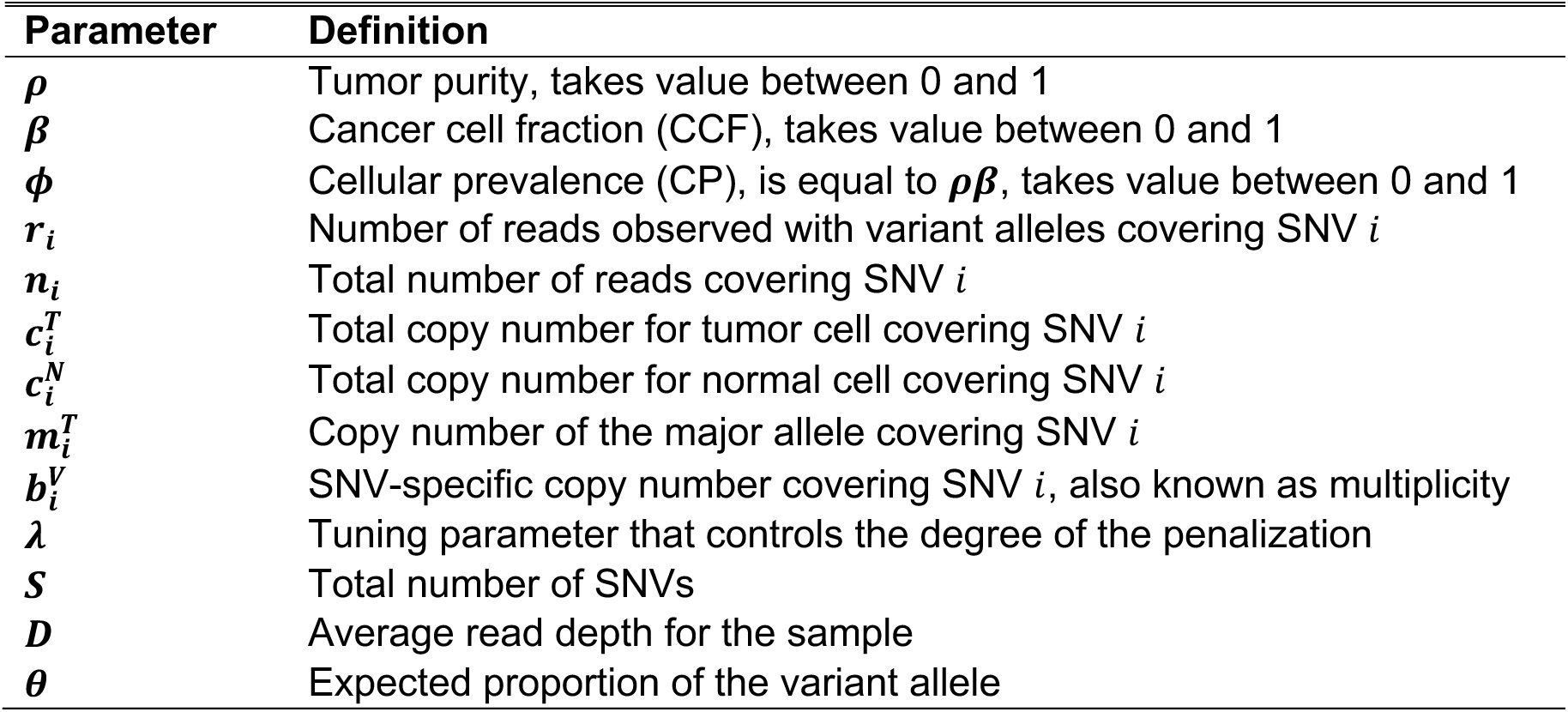

| Parameter | Definition |
| --- | --- |
| $\rho$ | Tumor purity, takes value between 0 and 1 |
| $\beta$ | Cancer cell fraction (CCF), takes value between 0 and 1 |
| $\phi$ | Cellular prevalence (CP), is equal to $\rho\beta$, takes value between 0 and 1 |
| $r_i$ | Number of reads observed with variant alleles covering SNV $i$ |
| $n_i$ | Total number of reads covering SNV $i$ |
| $c_i^T$ | Total copy number for tumor cell covering SNV $i$ |
| $c_i^N$ | Total copy number for normal cell covering SNV $i$ |
| $m_i^T$ | Copy number of the major allele covering SNV $i$ |
| $b_i^V$ | SNV-specific copy number covering SNV $i$, also known as multiplicity |
| $\lambda$ | Tuning parameter that controls the degree of the penalization |
| $S$ | Total number of SNVs |
| $D$ | Average read depth for the sample |
| $\theta$ | Expected proportion of the variant allele |

First, we assume that each sample is composed of two populations of cells with respect to the *i*-th SNV: one population containing normal cells or cancer cells without the SNV, and the other population of cancer cells harboring this SNV. We adopt the infinite sites assumption [39] to assume that there is a single origin for each SNV. Within cancer cells, SNV *i* may occur in one or a few cell populations (**Fig. 1a**), therefore we introduce *β_i_* to denote the cancer cell fraction (CCF) of SNV *i*, i.e., the fraction of cancer cells that carry SNV *i*. A clonal SNV exhibits a CCF of 1.0, indicating 100% presence in tumor cells. We further define the cellular prevalence (CP) of SNV *i* as *ϕ_i_*, so that *ϕ_i_* = *ρβ_i_*, where *ρ* denotes tumor purity, i.e., the proportion of cancer cells among all cells. Given *S* SNVs, we assume that there exist *K* groups of SNVs presenting distinct CCFs, where *K* is significantly smaller than *S*. We expect SNVs from a common cancer cell population or subclone to have identical CP (or CCF) and correspondingly share the same parameter *ϕ_i_* (or *β_i_*). We follow an established model for observed variant read counts [13], and assume that the observed number of variant reads *r_i_* at SNV *i* follows a binomial distribution *Binomial*(*n_i_*, *θ_i_*), where the total number of reads *n_i_* follow a Poisson distribution *Poisson*(*D*). We use 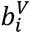 to denote the SNV-specific copy number at the *i*-th SNV. This measurement is distinct from allele-specific copy number when the SNV does not occur in all tumor cells. We then use 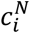 and 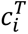 to denote the total copy number for normal and tumor cells, respectively.

We further express the expected proportion of the variant allele *θ_i_* as:

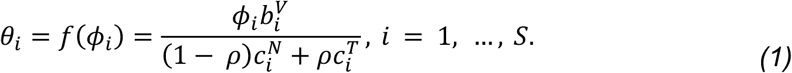

Hence the observed log-likelihood function, after canceling some constant, follows:

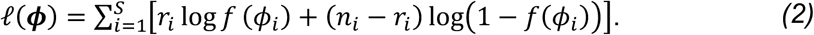

#### Input data

CliPP requires information of 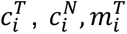, *r_i_*, *n_i_*, and *ρ* as inputs, which in order represent over SNV *i*: the total copy number for tumor cells, the total copy number for normal cells, the copy number of the major allele in tumor cells, the number of reads observed with variant alleles covering SNV, the total number of reads, and tumor purity. Parameters 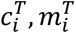, and *ρ* can be obtained using CNA-based deconvolution methods such as ASCAT [40], ABSOLUTE [41] and others [42–44], whereas 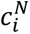 is typically set as 2. Parameter 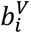, the SNV-specific copy number, is not directly observed or estimable using existing CNA software tools. Following a current convention that proved to be effective (see Eq 4 in Dentro et al. [35]), we assume 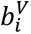 as:

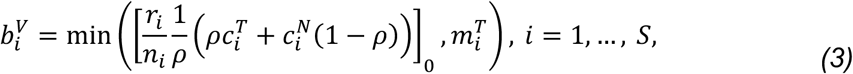

where [⋅]_0_rounds *x* to the nearest positive integer. Here, using 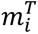 as the upper bound enforces the logical constraint that even if an SNV happened before any copy number event, the number of alleles carrying said SNV cannot exceed the total number of major alleles.

#### Output data

CliPP provides the inferred subclonal structure, including the total number of clusters, the total number of SNVs in each cluster, the estimated CP for each cluster, and the mutation assignment, i.e., cluster ID for each mutation. This output can then serve as the basis for various downstream analysis including the inference of phylogenetic trees [35].

### Parameter estimation using a regularized likelihood

The CPs, ***ϕ*** = (*ϕ*_1_, …, *ϕ_S_*)*^T^* may be estimated by maximizing the corresponding log-likelihood. However, our primary goal is to identify a homogeneous structure, i.e., clusters, of the CPs across all SNVs. Penalized estimation is a canonical tool to achieve homogeneity detection and parameter estimation simultaneously [45]. Therefore, we introduce a pairwise penalty to seek such homogeneity in ***ϕ***. To facilitate computation, we employ a normal approximation of the binomial random variable, i.e., 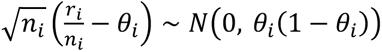, which leads to the following approximated log-likelihood

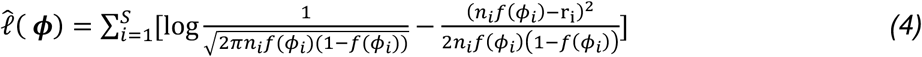

To promote homogeneity in ***ϕ***, we equip the negative log-likelihood −*l̂*(***ϕ***) with a pairwise shrinkage penalty, thereby solving the following optimization problem

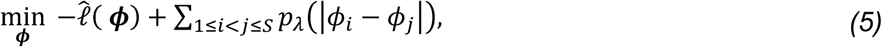

where *λ* > 0 is a tuning parameter that controls the degree of the penalization, and *p_λ_*(⋅) denotes a sparsity-inducing penalty function to identify the homogeneity structure in ***ϕ***, such as LASSO [46], SCAD [47], and MCP [48] to name a few. Here, we focus on the SCAD penalty which is defined as

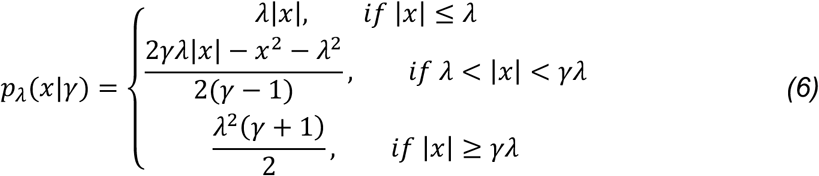

where *γ* is a hyper-parameter that controls the concavity of the SCAD penalty. Building upon the work of Fan and Li [47], we set *γ* to 3.7. These concave penalties offer sparsity similar to the LASSO penalty, allowing them to automatically yield sparse estimates. For variable selection in regression models, SCAD-penalized estimators can achieve the oracle property under suitable regularity conditions [47]. One can also choose other penalties, such as LASSO, and the formulation is straightforward. We have also implemented MCP as an alternative strategy.

Optimizing the objective function in Eq. (5) is nontrivial because the target variable ***ϕ*** is bounded, *i.e*., *ϕ_i_* = *ρβ_i_* ∈ [0,1], and the SCAD penalty is not convex. We employ a re-parametrization of *ϕ_i_* by defining 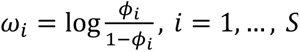, to remove the box constraint, which yields the following expression for 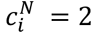.

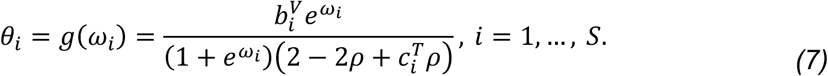

Note that *ω_i_*’s are monotonic with respect to *ϕ_i_*’s; *ω_i_* = *ω*_j_ implies *ϕ_i_* = *ϕ*_j_ and vice versa, thus the homogeneity pursuit of ***ϕ*** can be achieved by the homogeneity pursuit of ***ω***. We therefore perform a transformation on the loss function from Eq. (5), with the pairwise penalty applied to differences in ***ω*** to identify the homogeneity structure in ***ω***. Consequently, we formulate the optimization problem as follows:

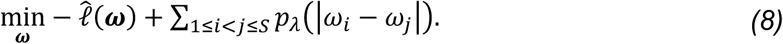

This is a non-convex optimization problem with respect to ***ω***, and its computation is non-trivial due to the complex log-likelihood. To address this challenge, we employ the alternating direction method of multipliers (ADMM) [49], where we approximate the negative log-likelihood at each iteration by a quadratic function, thereby simplifying the computation. Note that the SCAD penalty has zero derivative with respect to pairwise differences when (|*ω_i_* − *ω*_j_| > *γλ*). This property is particularly important in ADMM algorithms, as biases during the iterations could significantly impact the search for subgroups [50]. Further mathematical derivations and the computational construction of ADMM can be found in **Additional file 2: Section 1**.

Finally, it is crucial for the regularized likelihood-based approach to select a proper *λ*, in order to balance between over- and under-fitting. Here, the choice of *λ* determines the final number of clusters, with higher values yielding fewer clusters. Selection of *λ* therefore requires a balance between over-segmentation and excessive fusion. Although cross-validation and bootstrap-based selection are applicable in principle, they require repeated model fitting across candidate *λ* values and resampled datasets, substantially increasing computational cost. Standard AIC and BIC are also not straightforward to apply because CliPP uses a plug-in mutation-multiplicity estimate and an approximate likelihood, while the all-pairs SCAD penalty jointly determines cluster membership and cluster number, making the effective model degrees of freedom difficult to define. Instead, CliPP selects *λ* using a biologically motivated calibration criterion. In a bulk tumor sample, since the founding clonal mutations are expected to be present in all tumor cells, the highest inferred CP should approximate the tumor purity. Accordingly, for each fitted *λ*, we calculate the absolute relative difference 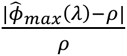, where *ϕ̂_max_*(*λ*) is the maximum cluster-level CP after post-processing. We retain *λ* values for which this relative difference is less than 1% and select the largest among them, because stronger penalization generally promotes greater pairwise fusion and yields fewer clusters. If no *λ* meets the 1% criterion, CliPP selects the *λ* that minimizes this relative difference, breaking ties in favor of the largest *λ*. In an empirical comparison, BIC-based selection was concentrated among smaller *λ* values, corresponding to less strongly fused solutions with more inferred clusters (**Additional file 1: Supplementary Fig. S1**). We therefore treated BIC-based selection as a sensitivity analysis rather than the primary model-selection criterion.

#### Post-processing of the clustering output

While the estimated CPs provide a natural definition of tumor subclonal architecture, manual post-processing is necessary in practice [13,51]. Penalized likelihood-based approach may, for example, produce spurious clusters containing a negligible number of SNVs. To reduce the need for manual curation, we apply the following post-filtering steps in sequence, with merging restricted to reconstructions containing more than two clusters: (1) Superclusters: when the maximum CP exceeds 1 and the corresponding cluster contains <40% of the sample’s SNVs, it is merged with the cluster with the next-highest CP. Such implausibly high estimates often reflect errors in the input CNA estimates. (2) Small clonal clusters: when the highest-CP cluster contains <15% of the sample’s SNVs, it is merged with the cluster with the next-highest CP. (3) Small subclonal clusters: clusters other than the highest-CP cluster that contain ≤5% of the sample’s SNVs are merged with their nearest neighbor in CP, starting with the eligible cluster with the lowest CP. Within each step, the merging condition is re-evaluated after each merge. The CP of a merged cluster is calculated as the SNV-count-weighted mean of the constituent cluster CPs. The current choice of cutoffs in these filters was trained with a sensitivity analysis using the PCAWG WGS data. These steps can be further modified by users when applying CliPP to a new dataset.

### Performance benchmarking using simulated datasets

We benchmarked CliPP against PhylogicNDT and PyClone-VI using two published cohorts, PhylogicSim500 and SimClone1000 [13], and our CliPPSim4k cohort. All 500 PhylogicSim500 tumors and 4,000 CliPPSim4k tumors were retained. A prespecified exclusion of SimClone1000 tumors with more than 50,000 SNVs before method fitting left 756 of the 965 published tumors. All methods were evaluated on the same eligible tumors, without post hoc sample exclusion based on reconstruction outputs.

PhylogicSim500 used PCAWG-derived copy-number profiles, with other parameters sampled independently from prespecified distributions, whereas SimClone1000 used a grid-based design with clonal copy-number states. The original benchmark roles and cohort characteristics are summarized in **Additional file 1: Supplementary Table S1**.

CliPPSim4k comprises 4,000 simulated tumors distributed across 27 combinations of mean sequencing depth *D* ∈ {100,200,500}, tumor purity *ρ* ∈ {0.4,0.6,0.9} and nominal CNA rate ∈ {0, 0.1, 0.2} (**Fig. 2a**). Each combination was assigned 148 tumors, with one additional tumor assigned to each of four design cells; all simulations used a fixed random seed. At the mutation-cluster level, the total SNV count *S* was sampled from a discrete uniform distribution over the integers 200-800, and a raw subclonal-mutation proportion from a continuous uniform distribution on [0,0.8). Proportions below 0.1 were set to zero, yielding fully clonal tumors (*K* = 1). Otherwise, the subclonal SNV count was obtained by multiplying *S* by the raw proportion and rounding down, and the total cluster number *K* was sampled uniformly from feasible values in {2,3,4}, subject to at least 20 SNVs per cluster. Subclonal SNVs were distributed as evenly as possible across the *K* − 1 subclonal clusters, with the remaining SNVs assigned to the clonal cluster. The realized subclonal-mutation proportion used as simulation truth was the subclonal SNV count divided by S.

**Figure 2:**
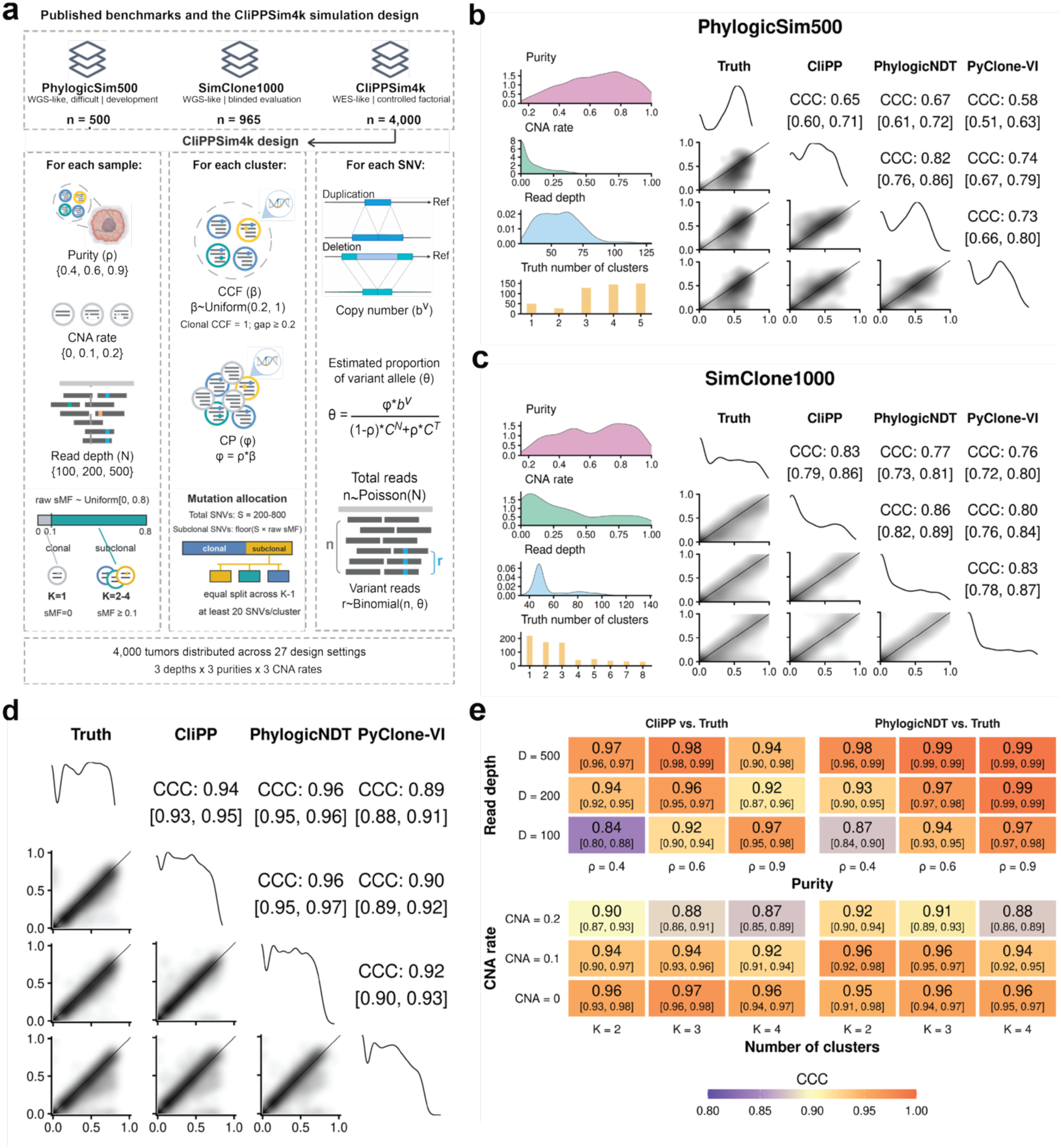
Simulation benchmarks for CliPP. **(a)** Overview of three simulation datasets and CliPPSim4k simulation design (see **Methods**). The two published WGS-like resources include PhylogicSim500 (n = 500), a development benchmark with disclosed ground truth [13], and SimClone1000 (published n = 965; analysis n = 756 after excluding tumors with more than 50,000 SNVs), an independent benchmark [13] originally used for blinded evaluation. Our CliPPSim4k comprises 4,000 WES-like tumors distributed across 27 combinations of mean read depth, tumor purity, and CNA rate; sample-, cluster-, and SNV-level simulation pipeline are shown. **(b&c)** Simulation characteristics and pairwise comparisons of the true and estimated proportions of mutations classified as subclonal in PhylogicSim500 and SimClone1000. Performances of three methods are shown: CliPP, PhylogicNDT and PyClone-VI (recommended setting: -r 10 restarts). Diagonal plots show marginal distributions, lower-triangle plots show pairwise distributions, and upper-triangle entries report CCC with 95% CIs. **(d)** Corresponding pairwise comparison in CliPPSim4k. **(e)** CCC for CliPP and PhylogicNDT within CliPPSim4k strata defined by read depth and purity (top) or CNA rate and true cluster number (bottom). Cluster-number-stratified analyses include tumors with 2 to 4 clusters.

The clonal cluster was assigned a CCF of 1; subclonal-cluster CCFs were sampled uniformly from [0.2,1) and redrawn until all cluster CCFs were separated by at least 0.2. Cellular prevalence was the product of cluster CCF and tumor purity. At the SNV level, each mutation was assigned independently to the copy-neutral or CNA-generating branch according to a Bernoulli distribution with the scenario-specific nominal CNA rate. SNVs in the copy-neutral branch had one mutant and one reference copy; in the CNA-generating branch, mutant and reference copy numbers were sampled uniformly from {1,2,3,4} and {0,1,2,3,4}, respectively. Expected VAFs *θ_i_* were calculated from tumor purity, cellular prevalence and SNV-specific copy number using Eq. (1). Total and variant read counts were generated as *n_i_* ∼ *Poisson*(*D*) and *r_i_* ∼ *Binomial*(*n_i_*, *θ_i_*), respectively. For each tumor, the simulator recorded purity, allele-specific copy-number states, variant and reference read counts, and mutation-level ground truth comprising cluster assignment, CCF and mutation multiplicity.

Agreement between estimated and true subclonal mutation fractions was assessed using Lin’s concordance correlation coefficient (CCC) [52], with 95% percentile confidence intervals from 1,000 tumor-level bootstrap resamples. One non-finite PhylogicNDT estimate in PhylogicSim500 was omitted only from CCC calculations involving that estimate. CliPPSim4k was evaluated overall and within prespecified read-depth-by-purity and CNA-rate-by-cluster-number strata. Tumors with *K* = 1 were omitted only from cluster-number-stratified CCC analyses because their true subclonal mutation fractions were uniformly zero.

Additional reconstruction accuracy was evaluated using three complementary metrics: the relative absolute error in cluster number (rdNC), the relative absolute error in clonal mutation fraction (rdCF), and the purity-normalized per-SNV CP root-mean-square error (RMSE):

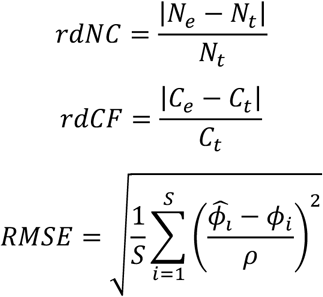

Total error was defined as the unweighted arithmetic mean of these three metrics:

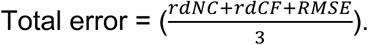

Here, *N_e_* and *N_t_* denote estimated and true cluster numbers, *C_e_* and *C_t_* denote estimated and true clonal mutation fractions, and *ϕ̂_i_* and *ϕ_i_* denote estimated and true cellular prevalences for SNV *i*. *S* is the SNV count and *ρ* is the true tumor purity. Normalization by *ρ* places CP errors on a common CCF scale across samples. Lower values indicate smaller errors; zero denotes exact agreement in the assessed quantity. For CliPPSim4k, total errors were compared between methods within strata defined separately by tumor purity, read depth, CNA rate or true cluster number, using paired, two-sided Wilcoxon signed-rank tests with Bonferroni correction across the three pairwise method comparisons in each stratum (**Additional file 1: Supplementary Tables S2-S4**).

### Subclonal reconstruction and benchmarking in PCAWG WGS tumors

The Pan-Cancer Analysis of Whole Genomes (PCAWG) study [13,53] dataset comprises whole-genome sequencing (WGS) data from 2,778 tumor samples obtained from 2,658 distinct donors across 39 cancer types, including 2,605 primary tumors and 173 metastases or recurrences (minimum average coverage at 30x in the tumor) [53]. PCAWG related data were downloaded from the ICGC data portal through the Bionimbus Protected Data Cloud (PDC). As input to CliPP, we use the consensus mutation calls derived from 4 distinct callers [53] and the consensus CNA calls from 6 distinct callers [13]. From these resources, per-SNV variant and total read counts, locus-specific total and major-allele copy numbers, and sample-level tumor purity were extracted and used as inputs to CliPP. The published PCAWG consensus subclonal assignments were retained solely as benchmark references and were not supplied to CliPP.

To benchmark CliPP in patient tumors, we compared its estimates with those from nine individual methods reported by the PCAWG study (BayClone [54], Ccube [55], cloneHD [56], CTPsingle [57], DPClust [26], PhylogicNDT [27], PhyloWGS [25], PyClone [24], Sclust [58]), two PyClone-VI [28] configurations, and the published PCAWG consensus [13]. PyClone-VI v0.2.0 was fitted with a maximum of 20 clusters and either 10 or 100 random restarts, denoted PyClone-VI1 and PyClone-VI2, respectively. After matching tumors across methods, 2,050 tumors had complete estimates across the comparison set. Application of the prespecified computational limit of 50,000 SNVs excluded 42 tumors, leaving 2,008 tumors for analysis.

Because the true subclonal architectures of patient tumors are not directly observable, the published PCAWG consensus was treated as an ensemble reference rather than ground truth. The comparison was based on the proportion of analyzed mutations estimated to be subclonal by each method. Pairwise agreement across tumors was quantified using CCC. For each method-versus-consensus comparison, 95% confidence intervals were obtained from 1,000 nonparametric bootstrap resamples of tumors. Pearson correlation coefficients, two-sided P values, and ordinary least-squares regression were reported.

### Computation benchmark

We measured the elapsed real time for each method from initiation to completion on the same machine equipped with Intel(R) Xeon(R) 6517P CPU @ 3.20GHz, using 16 CPU cores for consistency. In order to compute the speed for PhyloWGS, PyClone, and PhylogicNDT, which are time-consuming to run, we employed a random sampling approach on the PCAWG and TCGA datasets which gives us 111 samples from PCAWG and 87 samples from TCGA. After setting the minimum and maximum number of SNVs per sample to 20 and 20,000, respectively, we found that the SNV counts of the selected samples ranged from 24 to 14,284. PhyloWGS v1.0-rc2 was executed using the same settings applied in the simulation study. PyClone v0.13.1 was run with both recommended and default configurations (https://github.com/Roth-Lab/pyclone) [59]. PyClone-VI v0.2.0 was applied to all PCAWG samples using recommended and default settings, incorporating 20 clusters for fitting (-c 20) and either 10 (PyClone-VI1) or 100 restarts (PyClone-VI2) of variational inference (-r 10/100) (https://github.com/Roth-Lab/pyclone-vi) [60]. PhylogicNDT v1.0 (https://github.com/broadinstitute/PhylogicNDT) [61] was executed for 100 iterations (-ni 100), with the command to calculate ccf histograms (--maf_input_type calc_ccf). TCGA papillary thyroid carcinoma (PTC) WGS (n = 158) were evaluated using two CliPP execution modes, CPU and GPU (NVIDIA L40), with 10 repetitions per sample under the same benchmarking conditions.

### Concordance of driver mutation subclonal annotation with the PCAWG consensus

We compared per-mutation clonal/subclonal calls between CliPP and the PCAWG consensus subclonal reconstruction results [13] on the PCAWG ICGC sample set. Consensus calls and the corresponding driver-mutation annotations were obtained from the ICGC data repository and matched to CliPP calls by sample ID, chromosome, and genomic position. We report the overall concordance index between listed methods and consensus results.

### Number of reads per tumor chromosomal copy (NRPCC) as the quality indicator

Raw sequencing depth alone does not determine the reliability of cellular-prevalence estimation, because the informative variant signal is diluted by normal-cell contamination and by the number of tumor chromosomal copies over which reads are distributed. NRPCC [62] is used as an effective tumor-copy coverage metric for subclonal reconstruction. By adjusting expected total copy burden and tumor purity, NRPCC directly controls the reads for cellular prevalence estimations, and hence subclonal reconstruction. Low NRPCC reduces the separation between clonal and subclonal read count patterns relative to binomial sampling noise, making nearby CCF clusters less identifiable and increasing the instability of CP/CCF estimates. Conversely, higher NRPCC provides greater effective evidence for resolving distinct subclonal populations. We used NRPCC as a preprocessing criterion to exclude samples with insufficient effective coverage for reliable subclonal reconstruction. In adherence to established best practices [62], we calculate 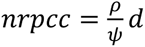, where *d* is the depth of sequencing, *ρ* is the purity and *ψ* is the average tumor sample ploidy, i.e., *ψ* = *ρψ_T_* + (1 − *ρ*)*ψ_N_*, where *ψ_T_* is tumor cell ploidy and *ψ_N_* = 2 is normal cell ploidy.

### Subclonal reconstruction in TCGA

The Cancer Genome Atlas (TCGA) contains whole-exome sequencing (WES) data from a cohort of tumor samples spanning 32 TCGA cancer types. The TCGA dataset consists of 9,815 WES samples with consensus mutation calls from 5 variant callers and 2 indel callers [63] with matched copy number segments and tumor purity estimates from ASCAT [40]. We implemented multiple steps of quality control and filtering of the TCGA samples, which are summarized in **Additional file 1: Supplementary Fig. S2, Supplementary Table S5.** Samples were excluded if they met any of the following conditions: (1) either too few SNVs (< 20) — insufficient statistical power for clustering — or too many SNVs (> 20,000), indicating potential artifacts; (2) low reads per chromosomal copy (nrpcc < 10); (3) low confidence copy number reconstruction, defined as meeting any of the following: ASCAT purity ≥ 0.99 (unrealistically high), genome instability ≤ 0.01 (minimal CN alteration), ASCAT purity < 0.40 (CN/CCF estimates unreliable), or low-event low-burden (fewer than 5 non-neutral copy-number segments AND total altered span < 40 Mb). After running CliPP, *s*amples were further excluded if (1) CliPP reported fewer than 4 SNVs assigned to the clonal cluster (insufficient clonal anchor for subclonal contrast); or (2) the largest inferred cluster CCF exceeded 1.2, indicating over-estimation of clonal CCF inconsistent with a biologically plausible solution. Silhouette-based cluster-separation quality control was applied to further remove samples with low quality subclonal reconstructions prior to biological analysis. For each multi-cluster reconstruction, we calculated the silhouette coefficient using mutation-level CP estimates and the final CliPP cluster assignments [64]. For each mutation *i*, *a_i_* was the mean absolute CP distance to other mutations in the same cluster, and *b_i_* was the smallest mean absolute CP distance to mutations in any other cluster. The mutation-level silhouette coefficient was calculated as *s_i_* = (*b_i_* − *a_i_*)/*max*(*a_i_*, *b_i_*), and the sample-level score was the mean of *s_i_* across mutations. Samples with a mean silhouette score < 0.4 were excluded from the high-confidence cohort. Single-cluster reconstructions, for which the silhouette coefficient is undefined, remained eligible if they satisfied all other quality-control criteria. Cancer types in which more than 25% of evaluable multi-cluster samples failed the silhouette criterion were excluded from cancer-type-level analyses; this study-specific rule excluded pancreatic cancer. The silhouette criterion was not applied to the simulation or PCAWG method-comparison benchmarks. The resulting high-confidence cohort comprised 5,424 samples across 31 cancer types.

### TCGA driver mutation annotation

Driver mutations were obtained from the IntOGen [65] driver annotation pipeline, and drivers identified in a given TCGA cancer type were extracted for 31 cancer types. After merging with the samples that passed CliPP filtering criteria, the resulting driver annotation set consisted of 4,444 samples with subclonal estimates for 14,797 SNV driver mutations in 285 unique genes.

#### Alignment-related QC

Sequence similarity can lead to ambiguous alignment and distort variant allele fractions, potentially producing false subclonal calls. To quantify mapping ambiguity at recurrent driver loci, we computed mapping quality scores for genes with at least 20 annotated driver mutations. For each genome position, we generated perfect 100-base read fastq files for every genome position and aligned them with BWA-mem [66], then extracted mapping quality scores (0-60) from the resulting BAM files. We then compare the distribution of mapping quality scores between clonal and subclonal mutations for the genes containing annotated mutation sites with a mapping quality score less than 60 using two-sided Wilcoxon rank sum tests. We found evidence of lower mapping quality scores among subclonal mutations in three of these genes (ATM, KMT2C and NCOR1; BH-adjusted [67] Wilcoxon p < 0.05; **Additional file 1: Supplementary Fig. S3**). To ensure high confidence in subclonality assignment in all downstream analyses, we removed mutations falling in regions with mapping quality score < 50; this filter eliminated the lower-confidence subset of *KMT2C* sites (107 of 339 mutations removed; 232 high-confidence sites retained). The final analysis set comprised 13,960 annotated driver mutations in 4,375 samples.

#### Pairwise driver-gene clonal timing

For each pair of distinct driver genes co-mutated in the same sample, the joint (A clonality, B clonality) call was classified as one of four direction categories — Forward (A clonal, B subclonal — A is likely acquired before B), Reverse (A subclonal, B clonal), Both clonal (no temporal information available from that sample), or Both subclonal (no temporal information). For the pan-cancer analysis, driver pairs were retained if at least ten tumors were resolvable (resolvable = Forward + Reverse). For cancer-type-specific analysis, a driver pair was required to have at least three resolvable tumors, with resolvable tumors comprising at least 20% of all tumors carrying that pair.

### CliPP-on-Web shiny application

To make subclonal reconstruction available to users without use of the command line, we developed CliPP-on-Web (https://bioinformatics.mdanderson.org/apps/CliPP/) [34], a web application built on CliPP 1.4.0 using R shiny (shiny 1.12.1, R 4.5.1) and deployed using Posit Connect. It takes the same inputs as the command line CliPP implementation but allows users to upload files using a simple graphical user interface. Subclonal reconstruction results are then visualized with interactive plots; and subclonal structure, mutation assignments, and plots can be downloaded per sample. See **Additional file 2: Section 2** for detailed advancement in CliPP 1.4.0/1.4.1. The shiny application also hosts download links for CliPP Sim4k simulation data, TCGA and PCAWG subclonal reconstruction results, along with driver clonality annotations for TCGA.

## Results

### CliPP reconstructs subclonal architecture through pairwise penalization

For a given tumor sample, mutation-level cellular-prevalence estimates may form a multimodal distribution, with distinct modes representing groups of mutations carried by clonal and subclonal cancer cell populations (**Fig. 1a-b**). CliPP estimates the cellular prevalence of each somatic single-nucleotide variant (SNV) from its variant and total read counts while accounting for tumor purity, local copy-number state and mutation multiplicity. Subclonal reconstruction can therefore be formulated as the identification of mutation clusters with homogeneous cellular-prevalence estimates, which together provide a quantitative representation of the tumor’s clonal and subclonal architecture.

CliPP implements this task through a standardized workflow that integrates somatic SNV read counts, allele-specific copy-number profiles and tumor purity (**Fig. 1c, Methods**). The input data are harmonized by mapping SNVs to copy-number segments and applying coverage and quality-control filters before mutation clustering. The fitted solution is then post-processed to merge biologically implausible or weakly supported clusters. For each sample, CliPP reports the inferred number of mutation clusters, mutation-to-cluster assignments, cluster sizes and cluster-level cellular-prevalence estimates, from which cancer cell fractions can be derived using tumor purity. These outputs support downstream evolutionary analyses and cohort-level characterization of clonal and subclonal architecture.

At the core of this workflow, CliPP performs homogeneity pursuit within a regularized maximum-likelihood framework (**Fig. 1d, Methods**). Conceptually, CliPP draws on homogeneity-pursuit methods such as fused LASSO and CARDS [37,50,68], but applies penalization to all pairwise differences among mutation-level cellular-prevalence parameters rather than requiring a prespecified ordering of mutations. The likelihood relates the observed variant and total read counts to mutation-level cellular prevalence while accounting for tumor purity, local copy-number state and mutation multiplicity. An all-pairs smoothly clipped absolute deviation (SCAD) fusion penalty [47] sets selected differences between cellular-prevalence estimates to zero, thereby assigning mutations with a common fitted cellular prevalence to the same cluster. Consequently, both the number of clusters and their membership emerge from the fitted model rather than being specified in advance.

CliPP fits the model over a sequence of penalty-parameter values λ, producing the solution path shown in **Fig. 1e**. As λ increases, progressively more mutation-level cellular-prevalence estimates are fused and fewer clusters are generally retained. The vertical dotted line in **Fig. 1e** marks the selected *λ*, at which the fused estimates define the final mutation clusters. CliPP obtains the solution path using an alternating direction method of multipliers algorithm (ADMM) [49] and selects the final solution by combining a clonal-anchor criterion with model parsimony (**Methods; Additional file 2: Section 1**). By casting cluster inference as a penalized optimization problem, CliPP achieves substantial computational gains while maintaining competitive reconstruction accuracy.

To facilitate use without local software installation, we also developed CliPP-on-Web, an interactive interface for running CliPP, visualizing reconstructed subclonal architectures and exploring the cohort-level results generated by CliPP.

### CliPP demonstrates high accuracy under simulation settings

Because ground-truth subclonal architectures are not directly observed in patient tumors, we benchmarked CliPP across three complementary simulation cohorts comprising 5,256 tumors (**Fig. 2a**). The two published WGS-oriented resources, PhylogicSim500 (n = 500) and SimClone1000 (n = 756 analyzed), were originally developed for method refinement, the former as training data and the latter for blinded evaluation, respectively [13]. We additionally generated CliPPSim4k (n = 4,000) to evaluate performance under controlled WES-like conditions. Together, these cohorts span broad variation in tumor purity, copy-number alterations, sequencing depth and the number of mutation clusters. Cohort definitions, inclusion criteria and simulation procedures are described in **Methods**; characteristics of the published benchmarks are summarized in **Additional file 1: Supplementary Table 1**.

We used the subclonal fraction, a measure of the proportion of analyzed mutations classified as subclonal, as the primary endpoint. This metric is focused on the dichotomization of clonal and subclonal mutations, representing the simplest, hence easiest yet biologically meaningful task in subclonal reconstruction. Agreement between estimated and true subclonal fraction was quantified using the Lin’s concordance correlation coefficient (CCC). A perfect CCC is 1. Using all finite estimate–truth pairs, CCC values in PhylogicSim500 were 0.65 (1,000 bootstrap 95% CI: 0.60, 0.71) for CliPP, 0.67 (95% CI: 0.61, 0.72) for PhylogicNDT, and 0.58 for PyClone-VI (95% CI: 0.51, 0.63, using the faster PyClone-VI1 with 10 restarts), with CliPP never trained from this data set (**Fig. 2b**). In SimClone1000 benchmark, the corresponding values were improved to 0.83 (95% CI: 0.79, 0.86), 0.77 (95% CI: 0.73, 0.81), and 0.76 (95% CI: 0.72, 0.80) (**Fig. 2c**). These external analyses therefore show that CliPP maintained competitive subclonal fraction accuracy across both published WGS-like resources with different roles and simulation designs. This also shows room for improvement for current methods on the simple task of annotating subclonal mutations at a generally low read depth.

Having established competitive performance to existing methods on the published resources, we next used CliPPSim4k to evaluate accuracy across a controlled WES-like design. All 4,000 simulated tumors were included. Across the complete cohort, the CCC between estimated and true subclonal fraction was 0.94 for CliPP (95% CI, 0.93, 0.95), 0.96 for PhylogicNDT (95% CI, 0.95, 0.96), and 0.89 for PyClone-VI (95% CI, 0.88, 0.91; **Fig. 2d**). PhylogicNDT demonstrated an overall best performance, with CliPP being a close second and competitive performance. CliPP concordance remained high across the factorial design: over the nine read-depth-by-purity settings, CCC values ranged from 0.84 to 0.98 for CliPP (**Fig. 2e**). The most challenging depth-purity setting for all three methods was mean read depth 100 and tumor purity 0.4, where CCC values were 0.84 (95% CI, 0.80, 0.88) for CliPP, 0.87 (95% CI, 0.84, 0.90) for PhylogicNDT, and 0.70 (95% CI, 0.64, 0.75) for PyClone-VI (**Additional file 1: Supplementary Table S2**). This shared decline indicates that limited signal-to-noise ratio (SNR) constrains subclonal fraction recovery across distinct modeling frameworks, although the reduction in concordance was most pronounced for PyClone-VI. Stratified CCC was not calculated for single-cluster tumors because their true subclonal fraction was invariant at zero.

To evaluate aspects of the reconstructed architecture not captured by subclonal fraction alone, we additionally assessed total reconstruction error, which sums up errors in the inferred number of clusters, subclonal fraction, and per-mutation cellular prevalence (**Methods**). Error distributions overlapped substantially across methods, although their relative ordering varied across simulation factors. When all three aspects were considered equally important, PhylogicNDT achieved the lowest error scores across three simulation datasets, with CliPP and Pyclone-VI following closely on all scenarios. Component-wise and total-error summaries for PhylogicSim500 and SimClone1000 are provided in **Additional file 1: Supplementary Table S3**, and for CliPPSim4k is provided in **Additional file 1: Supplementary Table S4**. Finally, reconstruction results by CliPP were broadly consistent when the SCAD penalty was replaced with the minimax concave penalty (MCP) in a sensitivity analysis (**Methods**, **Additional file 1: Supplementary Fig. S4**). Together, these benchmarks demonstrate that CliPP achieves robust subclonal fraction concordance and competitive reconstruction accuracy across diverse genomic data scenarios.

### CliPP achieves high concordance and enables rapid cohort-scale analysis in patient tumors

To evaluate CliPP performance in patient samples, we applied CliPP to WGS data from tumors in the Pan-Cancer Analysis of Whole Genomes (PCAWG) study, for which consensus subclonal reconstructions from 11 methods [13,28] are available. Using the fraction of subclonal mutations as the primary summary metric, CliPP achieved a close to perfect CCC of 0.93 (95% CI: 0.91, 0.95) with the PCAWG consensus **(Fig. 3a; Methods)**, presenting a similarly strong performance to PhylogicNDT’s 0.95 (95% CI: 0.93, 0.98), and significantly higher than all other evaluated methods. Pyclone-VI2 (slower Pyclone-VI with 100 restarts) showed only slightly improved concordance than PyClone-VI1. This suggests that CliPP’s optimization-based framework can achieve high accuracy comparable to state-of-the-art Bayesian approaches. In addition to calculating the CCC, we performed linear regression of the subclonal fractions estimated by the consensus versus each of the six top ranked methods across all samples (n=2,008): PhylogicNDT, CliPP, PyClone-VI1, Pyclone-VI2, Pyclone, PhyloWGS (**Fig. 3b**), and found a close to 1 coefficient for both PhylogicNDT (0.96) and CliPP (0.99), supporting their high concordance with consensus.

**Figure 3:**
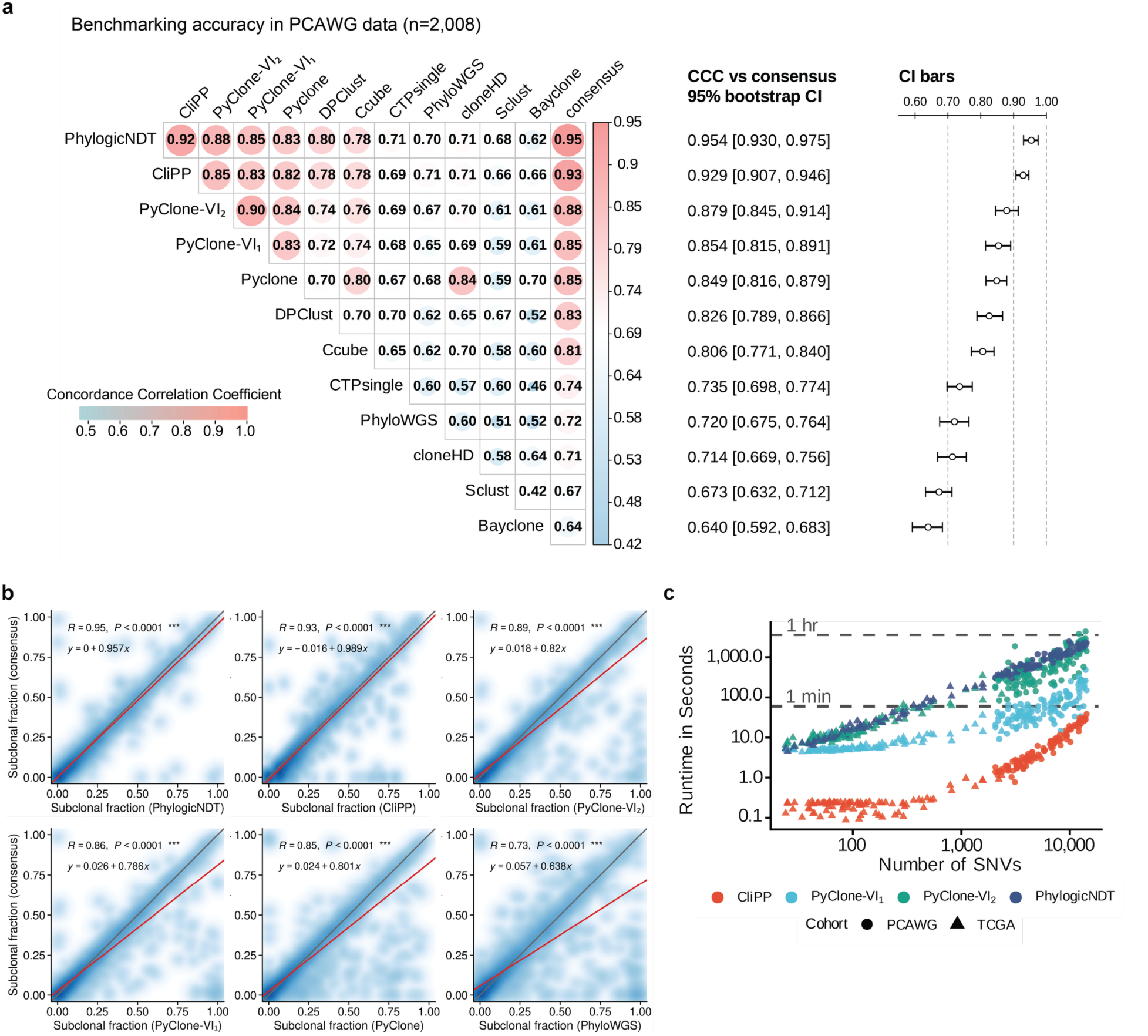
Concordance with the PCAWG consensus and computational performance in patient tumor data. **(a)** Pairwise Lin’s concordance correlation coefficients (CCCs) for the proportion of analyzed mutations classified as subclonal, estimated using CliPP, eight individual methods reported by the PCAWG study, two PyClone-VI configurations, and the published PCAWG consensus across 2,008 PCAWG tumors. The forest plot shows the CCC between each method and the consensus, with 95% confidence intervals obtained from 1,000 tumor-level bootstrap resamples. PyClone-VI1 and PyClone-VI2 denote runs using 10 and 100 variational-inference restarts, respectively. **(b)** Smooth scatter plots comparing estimates from selected methods with the PCAWG consensus. Pearson’s correlation coefficient and the fitted linear-regression equation are shown; the red line represents the fitted regression and the grey line the identity relationship. **(c)** Computational time over the number of SNVs for CliPP, PyClone-VI, and PhylogicNDT using 87 randomly selected WES samples and 111 randomly selected WGS samples up to 15,000 SNVs.

Beyond accuracy, CliPP achieved cohort-scale runtimes that shift subclonal reconstruction from a computational bottleneck to a routine, reproducible analysis step. Under identical configurations, CliPP was consistently faster than PhylogicNDT [27] and both PyClone-VI [28] configurations, with the largest relative advantage in tumors containing 100-1,000 mutations (**Fig. 3c, Methods**). This regime is particularly relevant to exome cohorts, as 75% of the analyzed TCGA tumors contained fewer than 200 SNVs. In aggregate cohort runs, CliPP processed the complete PCAWG (n = 2,008) and TCGA (n = 7,223) analysis sets in approximately one hour, representing at least a 10x reduction in runtime relative to PyClone-VI and PhylogicNDT under the benchmarked settings.

Cohort-scale analyses with the MCMC-based PhyloWGS and PyClone workflows required weeks to months on the same hardware. These computational demands are particularly relevant when review of mutation calls, copy-number and purity estimates, quality-control decisions or fitted clusters necessitates repeated analysis under revised configurations. In a separate benchmark, we ran CliPP ten times on the same cohort of 158 papillary thyroid carcinoma (PTC) WGS samples from TCGA, using identical input files and parameter settings. CliPP completed a cohort run within 10 minutes on average on the same computing platform, with sample-level runtimes on the order of seconds (**Methods, Additional file 1: Supplementary Fig. S5)**. These runtimes make iterative quality control and sensitivity analyses practical within the same inference framework and can leave more time for manual curation and biological interpretation. Together with CliPP’s high concordance with the PCAWG consensus, this efficiency supports reproducible subclonal reconstruction at cohort scale.

Having established CliPP’s performance in simulations and PCAWG whole-genome data, we next applied it to TCGA whole-exome tumors to systematically characterize pan-cancer subclonal architecture and driver mutation clonality.

### CliPP maps pan-cancer driver clonality and relative timing

Across 31 TCGA cancer types with cancer-type-specific driver annotations available from IntOGen [65], 6,355 tumors met our inclusion criteria for CliPP analysis. Following reconstruction-level quality control, 5,424 tumors formed the high-confidence cohort used for downstream analyses **(Methods; Additional file 1: Supplementary Fig. S2)**. This large cohort extends prior pan-cancer studies (∼1,200 [14] and ∼2,700 [17] whole exome sequenced, and ∼2,700 whole genome sequenced tumors [13]) and complements recent large-scale efforts [32] in providing an updated portrait of the driver evolutionary timing landscape across and within tumor types. Driver mutations are genetic alterations that contribute to, or “drive”, the development and progression of cancer [3,69]. Due to their role in tumor initiation and the fitness advantage they confer, driver mutations tend to be clonal. However, driver mutations can also arise later in evolution within subclones and confer a selective advantage later in tumor evolution [12,13].

We first benchmarked CliPP’s clonal/subclonal driver assignments against those produced by the 11-method PCAWG consensus pipeline [13]. Across 6,317 driver mutations from 1,597 PCAWG samples with consensus driver annotations, CliPP and the PCAWG consensus agreed on 96.3% of classifications (Cohen’s κ = 0.81), including 96.6% of consensus-clonal mutations and 93.5% of consensus-subclonal mutations (**Fig. 4a**). This high concordance achieved by a single computationally efficient method (finished analyzing PCAWG data in < 1hour) against a consortial effort of 11-method consensus supports the use of CliPP as a standalone tool for driver clonality assessment at scale.

**Figure 4:**
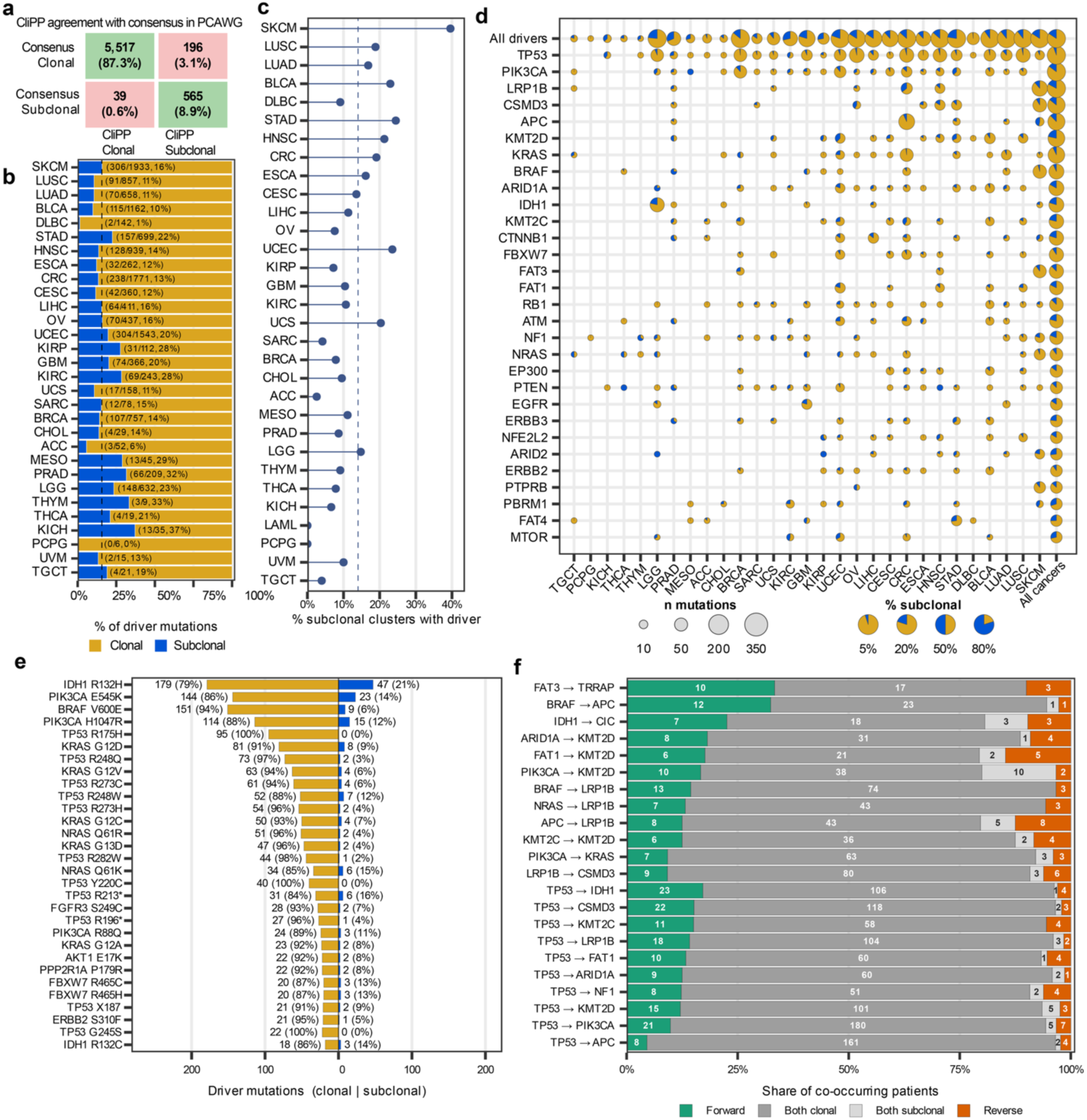
Subclonal driver mutation characterization across cancers with CliPP. **(a)** Comparison of subclonal assignment of driver genes in PCAWG between CliPP and the PCAWG consensus [13]. Cells show count and percent of driver mutations with clonality assignment (n = 6,317 paired driver calls). Overall agreement: 96.3%, Cohen’s κ = 0.81. **(b)** Proportion of driver mutations classified as subclonal in each cancer type in TCGA. Values next to each point indicate the number of subclonal driver mutations over the total number of driver mutations observed in a given cancer type (subclonal/total). Cancer types in **b-d** are ordered by median TMB. **(c)** Fraction of subclonal clusters that harbor at least one driver mutation by cancer type. Points show the percentage of subclonal clusters containing driver mutations. The dashed vertical line indicates the overall percentage across cancers (14%). (**d**) Clonal annotation of 30 most frequently mutated driver genes across cancer types. Each bubble is a scatter-pie whose area scales with the number of mutations in that (gene, cancer) cell and whose wedge split shows the clonal vs. subclonal share. Genes on the y-axis are ordered by total driver mutation count across cancers. **(e)** Clonal annotation of the 30 most frequently occurring driver mutations, ordered by the total mutation count. Mirrored butterfly bars show the clonal count (left, gold) and subclonal count (right, blue) for each locus; numerical labels at each bar tip give the count and percentage. **(f)** Pan-cancer pairwise driver-gene timing. For a pair labeled A → B, Forward indicates that A was clonal and B was subclonal; Reverse indicates that A was subclonal and B was clonal. For driver pairs with at least 10 tumors in which relative timing was resolvable (Forward or Reverse), all tumors harboring both drivers were included in the displayed distribution.

In TCGA, we use a robustly annotated set of high-confidence calls (**Additional file 1: Supplementary Figs. S2-S3**) covering 13,960 driver mutations in 281 genes across 4,375 patient samples [65]. We find that 2,189 (15.7%) driver mutations were subclonal (**Fig. 4b**). Among the cancer types profiled, chromophobe renal-cell carcinoma (KICH; 13/35, 37.1%), prostate adenocarcinoma (PRAD; 66/209, 31.6%) exhibit the highest proportion of subclonal driver mutations (**Fig. 4b**), while cutaneous melanoma (SKCM, n = 306) and endometrial carcinoma (UCEC, n = 304) contributed the largest raw count of subclonal driver mutations. Overall, driver mutations were reported in 14% of subclonal mutation clusters in TCGA, slightly higher than the 11% reported in the PCAWG WGS cohort [13] **(Fig. 4c**). Driver subclonality is broadly consistent across cancer types, with notable context-specific exceptions: although *TP53* was near-universally clonal, it showed an elevated subclonal fraction in prostate adenocarcinoma (PRAD, 27.6%), recapitulating the recurrent subclonal *TP53* mutations reported for prostate in the PCAWG cohort [13]. *LRP1B* was predominantly clonal in most cancers (17.1% subclonal pan-cancer; e.g. 11.2% in melanoma) yet was the largest single gene-level contributor to subclonal driver calls in colorectal cancer (38/107, 35.5% subclonal) (**Fig. 4d**).

At single-residue resolution, recurrent driver hotspots were almost predominantly clonal, with all 30 of the most frequently mutated hotspots having less than 25% subclonal presentation (**Fig. 4e**). The few subclonal-leaning exceptions fell in genes already enriched for subclonal calls, whereas *BRAF* p.V600E (n = 160, 5.6%), *KRAS* p.G12D (n = 89 9.0%) and the *TP53* DNA-binding-domain hotspots p.R175H (n = 95, 0%), p.R248Q (n = 75, 2.7%) and p.R273C (n = 65, 6.2%) were essentially fully clonal, consistent with strong selection for these specific substitutions at tumor initiation (per-cancer breakdowns in **Additional file 1: Supplementary Fig. S6**).

When multiple drivers are present within the same tumor, their clonal states provided coarse information about relative timing. If one driver was clonal and the other subclonal, the clonal driver was interpreted as having arisen earlier. In the pan-cancer analysis, the predominant pattern for pairs of drivers was co-clonal, accounting for 70–90% of co-mutated tumors for most pairs (**Fig. 4f**). These co-clonal classifications indicate that both events preceded the most recent common ancestor of the sampled tumor cells, and their relative order cannot be resolved from single-sample bulk sequencing. Among tumors in which the two drivers occupied different clonal states, the inferred orderings recapitulated established temporal sequences, both pan-cancer (**Fig. 4f**) and within individual cancer types (**Additional file 1: Supplementary Fig. S7**). Across cancers, *TP53* was resolved as the earlier driver in 17 out of 18 tissues in which it co-occurred, (the sole exception a 2-vs-2 tie in LIHC) consistent with its truncal role across cancers [12,13]. In clear cell renal cell carcinoma (KIRC), *VHL* was resolved to precede *SETD2* (**Additional file 1: Supplementary Fig. S7**), matching the established truncal *VHL*/3p-loss model [29,70]. Together, these results illustrate that CliPP’s subclonality calls capture biologically meaningful signal, and demonstrate the value of subclonal characterization in understanding driver mutation timing.

### CliPP-on-Web

We developed a web application, CliPP-on-Web, to allow researchers to perform subclonal reconstruction in a simple, user-friendly interface, which is available at https://bioinformatics.mdanderson.org/apps/CliPP [34] (**Fig. 5, Methods**). The CliPP web app leverages the computational advantages of CliPP to perform rapid subclonal reconstruction and interactive exploration of results. The CliPP web app was developed using CliPP (v1.4.0) as the back end, while the front end was built with R Shiny (v1.12.1) on R (v4.5.1). As illustrated in **Figure 5**, the CliPP web app enables users to run CliPP and visualize the distribution of VAF and estimated CP values across constructed subclones. This application will be a resource to allow quick exploration and annotation of subclonal mutations within user-samples, enabling further research into the evolutionary dynamics driving tumors.

**Figure 5:**
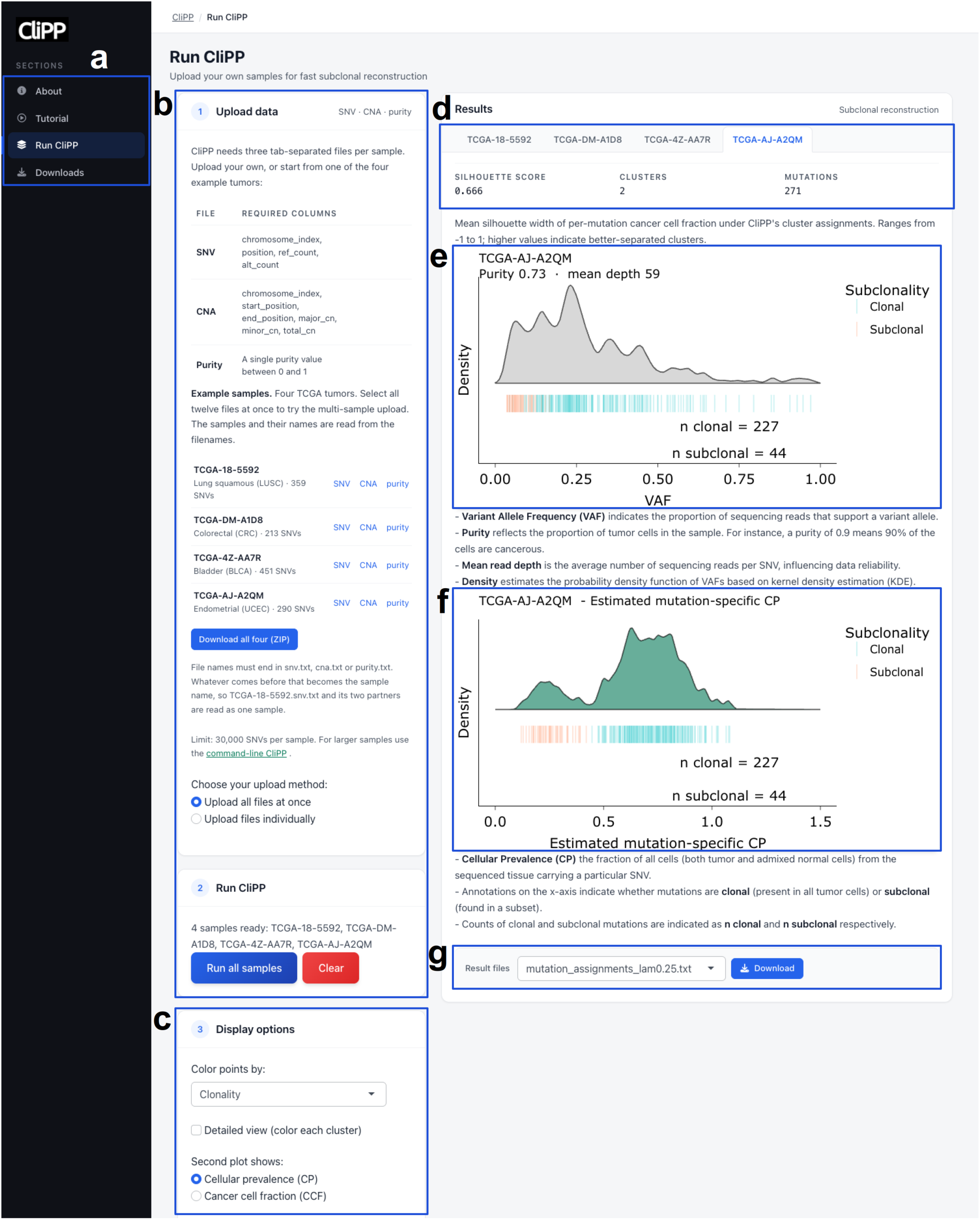
The CliPP-on-web Shiny application. **(a)** The navigation panel gives access to four sections: an overview and citation (“About”), an illustrated walkthrough (“Tutorial”), the analysis page shown here (“Run CliPP”), and precomputed CliPP output for TCGA and PCAWG cohorts, the, CliPPSim4k simulated cohort, and TCGA driver mutations annotated with clonality (“Downloads”). **(b)** CliPP requires three tab-separated files per sample: single nucleotide variants with read counts, copy-number segments, and a tumor purity estimate. **(c)** Display controls set point coloring (clonality or read depth), per-cluster coloring, the scale of the second plot (cellular prevalence or cancer cell fraction), and density smoothing. **(d)** Results are presented one tab per sample. For the selected sample, the silhouette score summarizes how cleanly the mutations separate, alongside the number of clusters and of clustered mutations. **(e-f)** Visualization of the VAF **(e)** and CP **(f)** distributions per sample. After performing subclonal reconstruction on a given sample(s), graphs showing the VAF and CP distribution are generated, with individual SNVs drawn as a “lawn” beneath each curve and colored by clonality (blue, clonal; red, subclonal). **(g)** Users can download the cluster assignments and subclonal structure of their samples. The CliPP-on-web Shiny app is available at https://bioinformatics.mdanderson.org/apps/CliPP/ [34].

## Discussion

We developed machine learning based CliPP for efficient subclonal reconstruction and used it to generate a quality-controlled pan-cancer map of tumor subclonal architecture and driver mutation clonality. Annotating clonal and subclonal mutations is a simpler task than building a full mutation phylogeny. It is a biologically useful task as demonstrated by our application to driver mutations. Yet this simple task is not entirely solved for single-sample DNA sequencing (DNAseq) data where the mutation-specific signal-to-noise ratio (SNR) remains low in highly complex regions of the cancer genomes. Single-sample DNAseq data are still more widely available than multi-sample, which has been shown to improve the SNR [10,11,70]. We utilized advanced optimization machine learning techniques to generate a subclonal reconstruction caller that is methodologically orthogonal to existing methods, in order to better support cancer researchers to analyze individual cohorts. CliPP can serve three purposes: 1) to achieve a better balance between accuracy and efficiency for calling subclonal mutations; 2) to provide an orthogonal method that captures a different set of signals, hence useful for consensus calling as needed; 3) to establish a new statistical framework that is more generalizable to multi-dimensions (multi-sample) than Bayesian methods.

Methodologically, CliPP reframes subclonal reconstruction as a constrained optimization problem rooted in penalized regression with a pairwise SCAD penalty [47], yielding sparse and homogeneous cellular-prevalence estimates and enabling fast, accurate clustering of mutations. Across >5,000 simulated samples, CliPP achieved competitive reconstruction accuracy to Bayesian approaches. In PCAWG real tumor samples, CliPP showed high concordance with consensus estimates of sample-level proportion of mutations classified as subclonal, while its driver-level clonal/subclonal assignments achieved 96.3% agreement with the previous consortial study. These complementary evaluations support its use as a standalone approach for cohort-scale reconstruction, while recognizing that the PCAWG consensus remains a reference rather than definitive ground truth. The substantial improvement in computational efficiency is important not only for initial cohort-wide analysis, but also for the repeated model fitting, input assessment, quality control and sensitivity analyses that are often required in practice. The standardized workflow further incorporates tumor purity, allele-specific copy number and NRPCC-based coverage assessment, addressing several input and quality-control factors known to influence subclonal reconstruction [62]. The CliPP-on-Web Shiny app (https://bioinformatics.mdanderson.org/apps/CliPP) [34] makes the method accessible to investigators without extensive computational expertise.

Several limitations should be considered. As with any single-sample sequencing analysis, distinct evolutionary histories (e.g., branching, punctuated, neutral) [62,71] can yield similar cluster configurations; thus, variability in subclonal architectures should be interpreted without assuming a specific evolutionary model. Accurate clonality calls depend on reliable purity and (clonal) copy-number inputs; errors in these inputs and unresolved subclonal CNAs can bias mutation-multiplicity and cellular-prevalence estimates. The current approximation on the unseen mutation multiplicity (Eq. 3) remains a major source of error for most subclonal callers. The current work centers on SNVs and small indels, and future efforts incorporating structural variants and subclonal CNA could provide a more complete view of tumor evolution. Future extension of the CliPP framework to multi-sample DNAseq is conceptually straightforward but will require the construction and comparison of multi-dimensional pairwise fusion techniques, improved estimation of mutation-multiplicity and a well-tested multi-dimensional hyperparameter selection strategy.

The large-scale driver-clonality maps and cohort-level reconstructions generated in our work provide a community resource that can be integrated with the extensive molecular, multi-omic and clinical annotations available through TCGA to support hypothesis generation and evaluation in independent cohorts. Together with the publicly accessible software and CliPP-on-Web application, these developments provide a flexible foundation for methodological and biological advances and may help connect tumor evolutionary analysis with precision oncology. Any clinical application will require appropriately designed studies and prospective validation.

## Conclusions

CliPP provides a scalable framework for reconstructing tumor subclonal architecture from bulk DNA sequencing data. By combining competitive reconstruction performance, substantially improved computational efficiency and a standardized quality-control workflow, CliPP enables reproducible application across large sequencing cohorts. Its pan-cancer application shows that most recurrent driver mutations are clonal, while a substantial minority arise subclonally in gene- and cancer-type-dependent patterns. CliPP and the accompanying cohort-level resources therefore provide a practical foundation for large-scale studies of tumor evolution and for future analyses using an aggregate of longitudinal, multi-region and single-cell data.

## Abbreviations

ADMM: Alternating direction method of multipliers
AIC: Akaike information criterion
ASCAT: Allele-specific copy-number analysis of tumors
BAM: Binary alignment map
BH: Benjamini-Hochberg
BIC: Bayesian information criterion
CARDS: Clustering algorithm in regression via data-driven segmentation
CCC: Concordance correlation coefficient
CCF: Cancer cell fraction
CI: Confidence interval
CliPP: Clonal structure identification through Pairwise Penalization
CN: Copy number
CNA: Copy-number alteration
CP: Cellular prevalence
CPU: Central processing unit
GPU: Graphics processing unit
ICGC: International Cancer Genome Consortium
LASSO: Least absolute shrinkage and selection operator
MCMC: Markov chain Monte Carlo
MCP: Minimax concave penalty
NRPCC: Number of reads per tumor chromosomal copy
PCAWG: Pan-Cancer Analysis of Whole Genomes
QC: Quality control
rdCF: Relative absolute error in clonal mutation fraction
rdNC: Relative absolute error in cluster number
RMSE: Root-mean-square error
SCAD: Smoothly clipped absolute deviation
SNV: Single-nucleotide variant
TCGA: The Cancer Genome Atlas
TMB: Tumor mutational burden
VAF: Variant allele fraction
WES: Whole-exome sequencing
WGS: Whole-genome sequencing

## Data availability

Raw read counts of bulk DNA sequencing data, clinical data and somatic mutations from 10,223 tumor samples across 32 TCGA cancer types are available for download from the Genomic Data Commons data portal (https://portal.gdc.cancer.gov/) [31]. The tumor purity and copy number profiling of TCGA samples were obtained using ASCAT v3 [40] and the results were released on https://github.com/VanLoo-lab/ascat/tree/master/ReleasedData [72].

PhylogicSim500 and SimClone1000 dataset can be downloaded from https://data.mendeley.com/datasets/compare/by4gbgr9gd [73].

The PCAWG dataset was obtained from the ICGC data portal: https://dcc.icgc.org/pcawg [74]. Specifically, the SNV sequencing reads were downloaded through https://dcc.icgc.org/repositories [75], while the CNA and purity data were accessed through https://dcc.icgc.org/releases/PCAWG/consensus_cnv [76]. Sample information, including mapping from ICGC identifiers to TCGA identifiers, was downloaded through the ICGC portal. The ICGC data portal was retired in June 2024; the PCAWG data remain available as legacy ICGC 25K data through ICGC ARGO (https://docs.icgc-argo.org/docs/data-access/icgc-25k-data) [77].

The previous subclonal reconstruction results from 11 methods and the consensus calls can be downloaded from Zenodo at https://doi.org/10.5281/zenodo.14510686 [78].

Driver mutation annotations were downloaded from IntOGen at https://www.intogen.org/download?file=IntOGen-Drivers-20240920.zip [79]

Subclonal structure results of all samples and the identified subclonality of driver mutations for this study are available for download at https://bioinformatics.mdanderson.org/apps/CliPP/ [34]. Alternatively, CliPP data for CliPPSim4k, TCGA, and PCAWG can be accessed on Zenodo at https://doi.org/10.5281/zenodo.23424567 [80].

All other relevant data are available from the corresponding author upon reasonable request.

## Code availability

The CliPP model used for reconstructing tumor subclonal structure is freely available and can be downloaded from https://github.com/wwylab/CliPP [81]. CliPP versions 1.4.0/1.4.1 were used to generate the results in this work. Additionally, the code for the Shiny app can be found at https://github.com/wwylab/CliPP-ShinyApp.

## Funding

Y.J. is supported in part by RP210006. Y.J., M.D.M., Y.D., S.J. and W.W. are supported by R01CA268380. Q.T. and W.W. are supported by DoD PC210079. Q.T. is supported in part by R01CA275990. R.L. is supported in part by R01CA283402. T.L. and P.V.L. were supported by the Francis Crick Institute, which receives its core funding from Cancer Research UK (CC2008), the UK Medical Research Council (CC2008) and the Wellcome Trust (CC2008). P.V.L. is a CPRIT Scholar in Cancer Research and acknowledges CPRIT grant support (RR210006). S.J.S. was supported by the National Research Foundation of Korea (NRF) grant funded by the Korea government (MSIT), grant number 2022M3J6A1063595, and 2023R1A2C1006587. J.R.W. was supported by the Mark Foundation for Cancer Research ASPIRE award. M.K. is supported in part by R01CA268380.

## Acknowledgements

The results presented here are in part based upon data generated by the TCGA Research Network: https://www.cancer.gov/tcga [82]. We thank John Weinstein for his valuable feedback on this manuscript. Special thanks to Sylvain Laroche and Chris Wakefield from the Quantitative Research Computing team for their assistance in deploying the CliPP-on-Web and enabling public access. The author(s) acknowledge the support of the High Performance Computing for research facility at the University of Texas MD Anderson Cancer Center for providing computational resources that have contributed to the research results reported in this paper.

## Declaration of interests

Y.J. is currently a Manager, Statistics at AbbVie. The other authors declare no competing interests.

## Ethics approval and consent to participate

The study is conducted in accordance with the Declaration of Helsinki and approved by the Institutional Review Board of University of Texas, MD Anderson Cancer Center (IRB protocol PA14-0404 under PI Dr. Wenyi Wang). No patient was recruited for this study. Clinical trial number: not applicable.

## Consent for publication

This study used published data from patient samples. Patient consent is not applicable.

## Authors’ contributions

S.J.S. and W.W. conceived the project. Y.J., K.Y., S.J.S., H.Z. and W.W. developed the mathematical method. Y.J., Y.D. and K.Y. built the software tool. M.D.M., A.W. and Q.T. built and deployed the Shiny app. Y.J., M.D.M. and Y.D. performed the data analysis and interpretation, planned the figure design, and wrote the manuscript collaborating with all the other authors. S.J., Q.T. and S.C. assisted on data analysis and interpretation. R.L., Q.T. and X.L. assisted with data maintenance and manuscript writing. Y.T. contributed to the code optimization. T.L. assisted with curating the copy number data. J.R.W. advised on the data analysis. M.T. and P.V.L. assisted in data interpretation, model development and copy number profiling. M.K., X.L. and H.Z. advised on mathematical modeling. W.W. conceived the study, planned and supervised the work, performed the analysis, and wrote the manuscript collaborating with all the other authors. All authors contributed to the interpretation of results and commented on and approved the final manuscript.

## Notes

https://github.com/wwylab/CliPP

https://doi.org/10.5281/zenodo.23424567

### Summary of Updates

Updated results for CliPP version 1.4, with updated timing, accuracy benchmarks, and driver analysis.

