## Additional File 1 for "Scalable subclonal reconstruction of cancer cells in DNA sequencing data using a penalized likelihood model"

### **Additional file 1: Supplementary Tables and Figures**

### **Supplementary Tables**

| **Aspect** | **PhylogicSim500** | **SimClone1000** |
| --- | --- | --- |
| Primary purpose | Community benchmark simulation set for single-sample subclonal reconstruction | |
| Sample count | 500 simulated tumors | 965 published tumors; 756 analyzed after excluding tumors with >50,000 SNVs. |
| How scenarios are sampled | Copy-number profiles sampled from real PCAWG tumors; other parameters sampled independently from fixed distributions | Grid design covering a broad set of scenario combinations encountered in PCAWG |
| Ground truth availability | Ground-truth architectures were released with the data to allow debugging/tuning | Ground truth not released before final evaluation; design kept hidden to make reverse-engineering difficult |
| Copy-number complexity note | Implicitly built from PCAWG CN profiles; the simulation description emphasizes realistic CN backgrounds | Clonal copy-number states; subclonal CNAs were not simulated. |

**Supplementary Table S1. Characteristics of the published simulation benchmarks and analysis cohorts.** The original resources were introduced by Dentro et al.[1]. In PhylogicSim500, tumor purity ranged from 0.16 to 0.99, CNA rate from 0 to 0.53, read depth from 27 to 129, and the true number of mutation clusters from 1 to 5. In SimClone1000, tumor purity ranged from 0.16 to 1.00, CNA rate from 0 to 1.00, read depth from 34 to 149, and the true number of mutation clusters from 1 to 8. CNA rate denotes the proportion of SNVs in copy-number-altered regions.

|  | **ρ = 0.4** | **ρ = 0.6** | **ρ = 0.9** |  | **K = 2** | **K = 3** | **K = 4** |
| --- | --- | --- | --- | --- | --- | --- | --- |
| **D = 500** | 0.93 [0.88, 0.97] | 0.88 [0.82, 0.93] | 0.91 [0.85, 0.94] | **CNA = 0.2** | 0.89 [0.83, 0.94] | 0.86 [0.82, 0.89] | 0.85 [0.82, 0.88] |
| **D = 200** | 0.93 [0.90, 0.96] | 0.91 [0.86, 0.95] | 0.95 [0.93, 0.98] | **CNA = 0.1** | 0.79 [0.71, 0.87] | 0.89 [0.85, 0.92] | 0.84 [0.81, 0.87] |
| **D = 100** | 0.70 [0.64, 0.75] | 0.91 [0.87, 0.94] | 0.94 [0.90, 0.97] | **CNA = 0** | 0.81 [0.73, 0.88] | 0.85 [0.80, 0.89] | 0.87 [0.84, 0.90] |

**Supplementary Table S2.** PycloneVI’s sMF vs. Truth CCC on CliPPSim4k under different settings.

|  |  | **SimClone**  **1000** | **PhylogicSim500** | **CliPPSim4k** |  | **SimClone**  **1000** | **PhylogicSim500** | **CliPPSim4k** |
| --- | --- | --- | --- | --- | --- | --- | --- | --- |
|  | **rdCF** |  |  |  | **rdNC** |  |  |  |
| **CliPP** |  | 0.454 ± 2.184 | 0.283 ± 0.309 | 0.109 ± 0.176 |  | 0.345 ± 0.555 | 0.306 ± 0.274 | 0.160 ± 0.323 |
| **PhylogicNDT** |  | 0.510 ± 2.261 | 0.264 ± 0.323 | 0.096 ± 0.153 |  | 0.234 ± 0.274 | 0.288 ± 0.246 | 0.100 ± 0.193 |
| **PyClone-VI** |  | 0.518 ± 2.406 | 0.318 ± 0.395 | 0.131 ± 0.307 |  | 0.242 ± 0.276 | 0.357 ± 0.241 | 0.131 ± 0.209 |
|  | **RMSE** |  |  |  | **Total error** |  |  |  |
| **CliPP** |  | 0.114 ± 0.078 | 0.153 ± 0.070 | 0.094 ± 0.056 |  | 0.304 ± 0.770 | 0.247 ± 0.169 | 0.121 ± 0.140 |
| **PhylogicNDT** |  | 0.099 ± 0.083 | 0.145 ± 0.070 | 0.091 ± 0.060 |  | 0.281 ± 0.784 | 0.232 ± 0.171 | 0.096 ± 0.098 |
| **PyClone-VI** |  | 0.107 ± 0.091 | 0.144 ± 0.071 | 0.085 ± 0.056 |  | 0.289 ± 0.831 | 0.273 ± 0.197 | 0.116 ± 0.148 |

**Supplementary Table S3.** Three methods’ mean ± standard deviation of rdCF, rdNC, RMSE, and total error from three simulation cohorts.

| **CliPPSim4k** | **n** | **CliPP** | **PhylogicNDT** | **PyClone-VI** |
| --- | --- | --- | --- | --- |
| Purity 40% | 1,335 | 0.159 ± 0.155 | 0.119 ± 0.114 | 0.138 ± 0.156 |
| Purity 60% | 1,332 | 0.114 ± 0.120 | 0.093 ± 0.089 | 0.110 ± 0.148 |
| Purity 90% | 1,333 | 0.090 ± 0.133 | 0.076 ± 0.084 | 0.099 ± 0.136 |
| CNA rate 0% | 1,332 | 0.097 ± 0.141 | 0.069 ± 0.092 | 0.102 ± 0.154 |
| CNA rate 10% | 1,336 | 0.123 ± 0.144 | 0.094 ± 0.097 | 0.119 ± 0.154 |
| CNA rate 20% | 1,332 | 0.143 ± 0.130 | 0.124 ± 0.098 | 0.126 ± 0.133 |
| Read depth 100 | 1,333 | 0.156 ± 0.153 | 0.124 ± 0.110 | 0.147 ± 0.159 |
| Read depth 200 | 1,334 | 0.118 ± 0.142 | 0.090 ± 0.091 | 0.095 ± 0.127 |
| Read depth 500 | 1,333 | 0.089 ± 0.113 | 0.073 ± 0.085 | 0.105 ± 0.149 |
| True clusters = 1 | 515 | 0.046 ± 0.167 | 0.002 ± 0.028 | 0.029 ± 0.099 |
| True clusters = 2 | 1,220 | 0.098 ± 0.138 | 0.092 ± 0.108 | 0.111 ± 0.177 |
| True clusters = 3 | 1,145 | 0.140 ± 0.135 | 0.106 ± 0.088 | 0.126 ± 0.134 |
| True clusters = 4 | 1,120 | 0.161 ± 0.111 | 0.131 ± 0.089 | 0.150 ± 0.127 |

**Supplementary Table S4.** Mean ± standard deviation of CliPPSim4k’s total errors stratified based on settings.

| Abbreviation | Cancer type | Abbreviation | Cancer type |
| --- | --- | --- | --- |
| ACC | Adrenocortical carcinoma | **LIHC** | Hepatocellular carcinomas |
| BLCA | Bladder urothelial carcinoma | **LUAD** | Lung adenocarcinoma |
| BOCA | Bone cancer | **LUSC** | Lung squamous cell carcinoma |
| BRCA | Breast invasive carcinoma | **MALY** | Malignant lymphoma |
| BTCA | Biliary tract cancer | **MESO** | Mesothelioma |
| CESC | Cervical squamous cell carcinoma & adenocarcinoma | **OV** | Ovarian serous cystadenocarcinoma |
| CHOL | Cholangiocarcinoma | **PAAD** | Pancreatic adenocarcinoma |
| CLLE | Chronic lymphocytic leukemia | **PBCA** | Pediatric brain cancer |
| CMDI | Chronic myeloid disorders | **PCPG** | Pheochromocytoma and paraganglioma |
| CRC | Colorectal carcinoma | **PRAD** | Prostate adenocarcinoma |
| DLBC | Diffuse large B-cell lymphoma | **SARC** | Sarcoma |
| ESCA | Esophageal carcinoma | **SKCM** | Melanoma |
| GBM | Glioblastoma multiforme | **STAD** | Stomach adenocarcinoma |
| HNSC | Head & neck squamous cell carcinoma | **TGCT** | Testicular germ cell tumors |
| KICH | Renal chromophobe | **THCA** | Thyroid papillary carcinoma |
| KIRC | Renal clear cell carcinoma | **THYM** | Thymoma |
| KIRP | Renal papillary carcinoma | **UCEC** | Uterine corpus endometrial carcinoma |
| LAML | Acute myeloid leukemia | **UCS** | Uterine carcinosarcoma |
| LGG | Brain lower grade glioma | **UVM** | Uveal melanoma |

**Supplementary Table S5.** Acronyms used for cancer types analyzed in this study.

**Supplementary Figures**

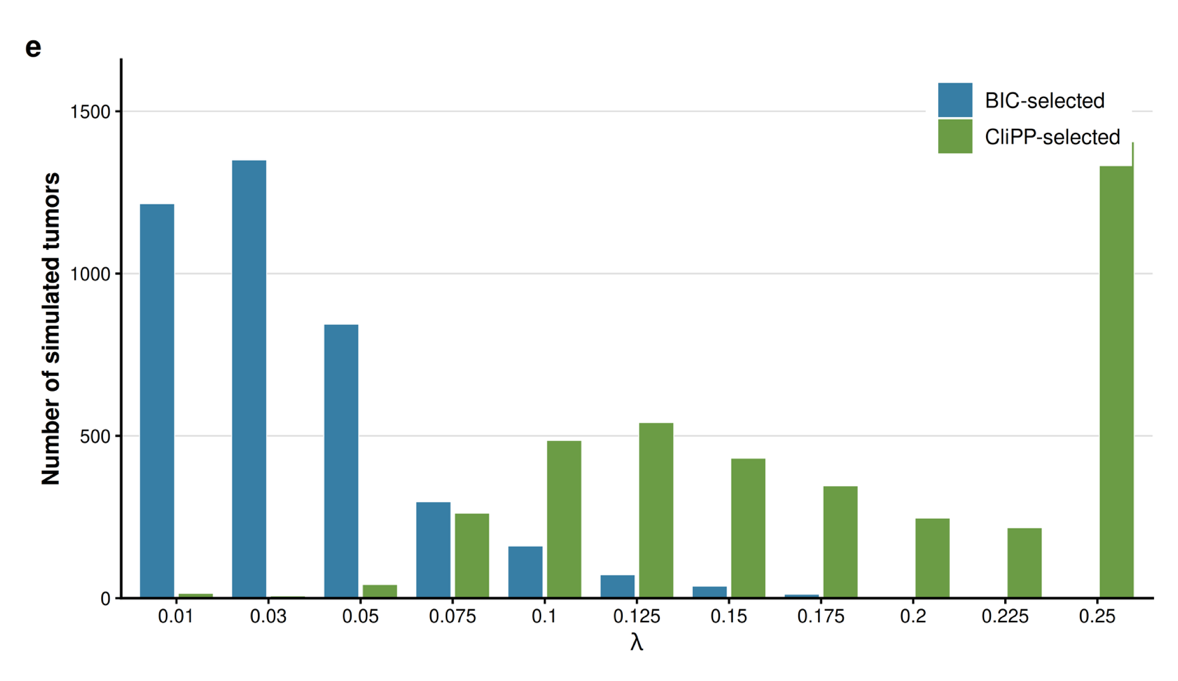

**Supplementary Figure S1:** Grouped bar chart showing comparisons between model selections based on BIC and proposed *ad hoc* approach in CliPPSim4k using the same
$\lambda$ sequence.

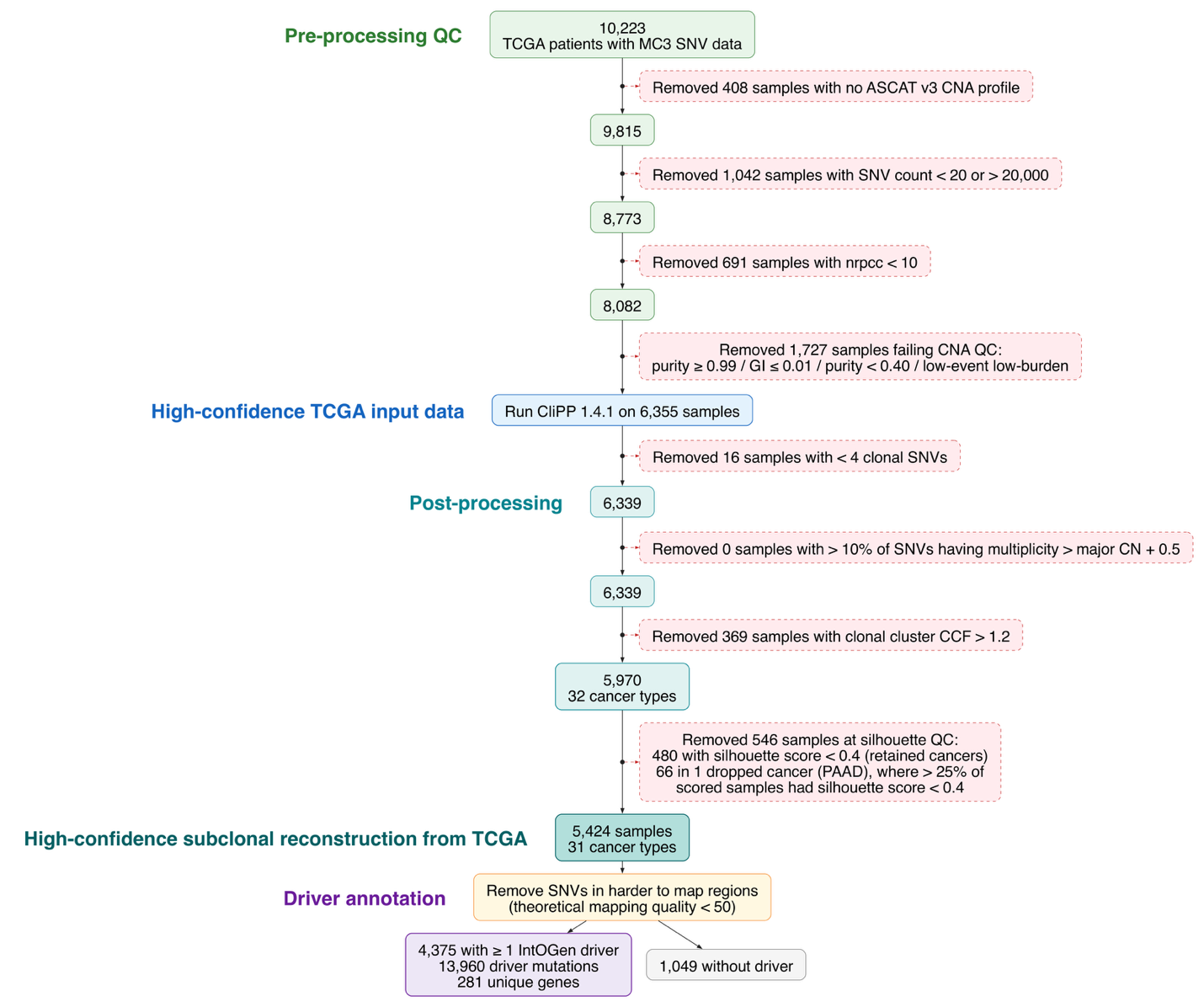

**Supplementary Figure S2: Overview of the TCGA data quality control filtering.** CONSORT diagram for data processing in TCGA.

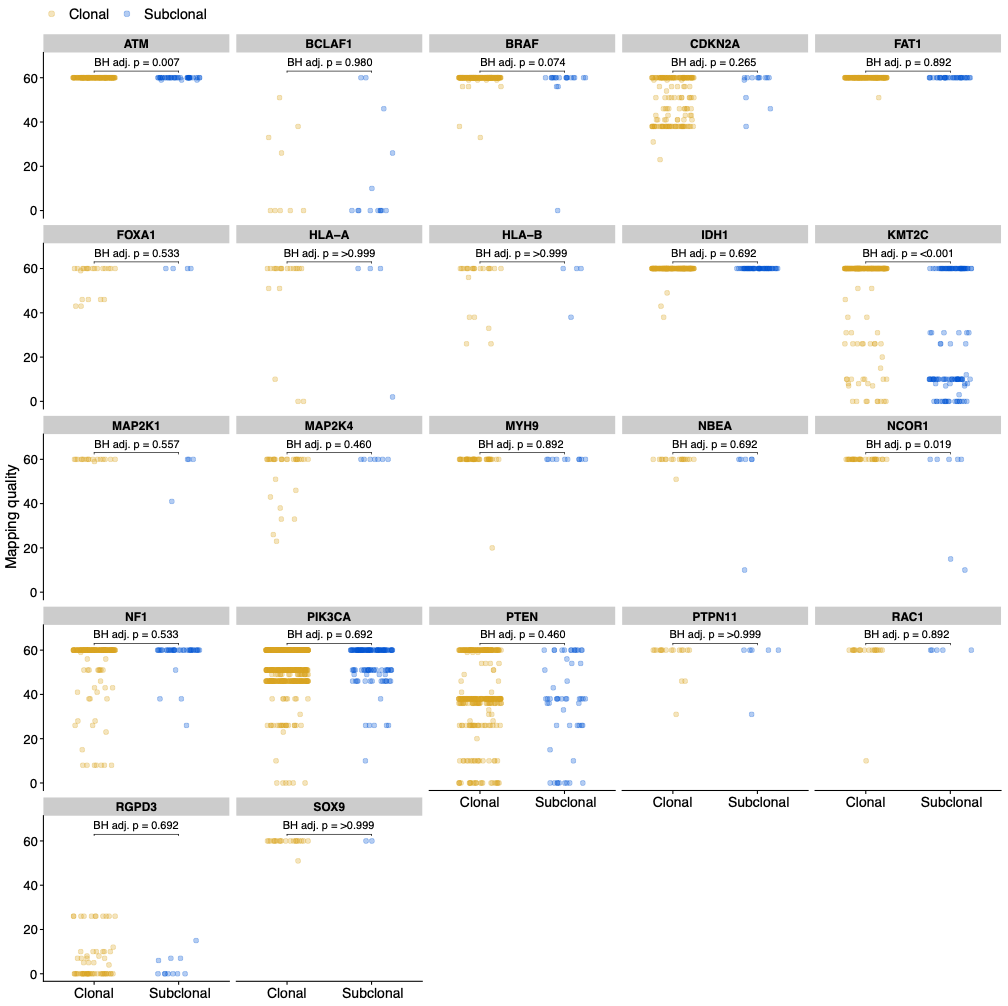

**Supplementary Figure S3: Assessment of potential mapping-related confounding in driver-mutation clonality assignments.**  Scatterplots of the mapping quality scores for all mutations in driver genes that have non-perfect mapped reads. Each point represents one mutation in one tumor. Within each gene, mapping-quality score distributions were compared between clonal (blue) and subclonal (yellow) mutations using two-sided Wilcoxon rank-sum tests; displayed P values were adjusted across genes using the Benjamini-Hochberg procedure.

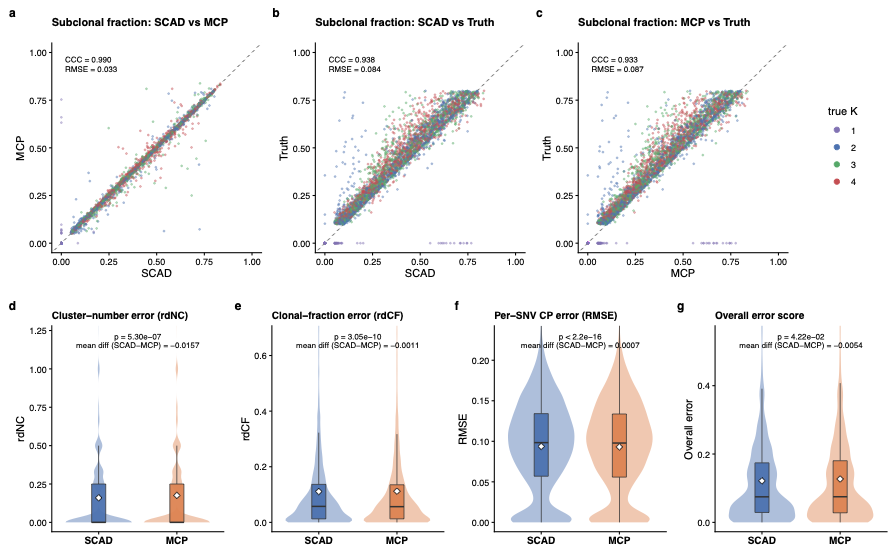

**Supplementary Figure S4: Comparison of SCAD and MCP penalties in CliPPSim4k (a-c)** Agreement in the estimated proportion of subclonal mutations between SCAD and MCP fits **(a)**, SCAD and simulation truth **(b)**, and MCP and simulation truth **(c)**. Each point represents one simulated tumor (n = 4,000) and is colored by the true number of clusters (K); dashed lines indicate equality. Lin’s concordance correlation coefficient (CCC) and root-mean-square error (RMSE) are shown. **(d-g)** Paired distributions of cluster-number error (rdNC; **d**), clonal-fraction error (rdCF; **e**), per-SNV cellular-prevalence RMSE **(f)** and overall error score (g). Lower values indicate better reconstruction. Boxes show medians and interquartile ranges, and white diamonds indicate means. P values are from two-sided Wilcoxon signed-rank tests.

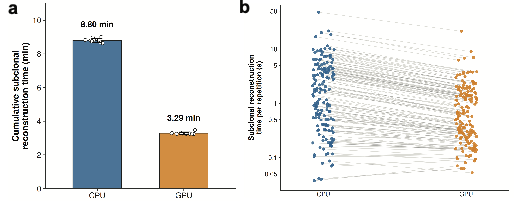

**Supplementary Figure S5. Speed summary of CliPP’s two modes on the TCGA papillary thyroid cancer whole-genomes (PTC WGS). (a)** Cumulative subclonal reconstruction runtime (minutes) for CliPP on TCGA PTC WGS (n = 158) using CPU and GPU execution modes. **(b)** Paired mean runtime (seconds) per sample for CPU (Intel(R) Xeon(R) 6517P) and GPU (NVIDIA L40) execution.

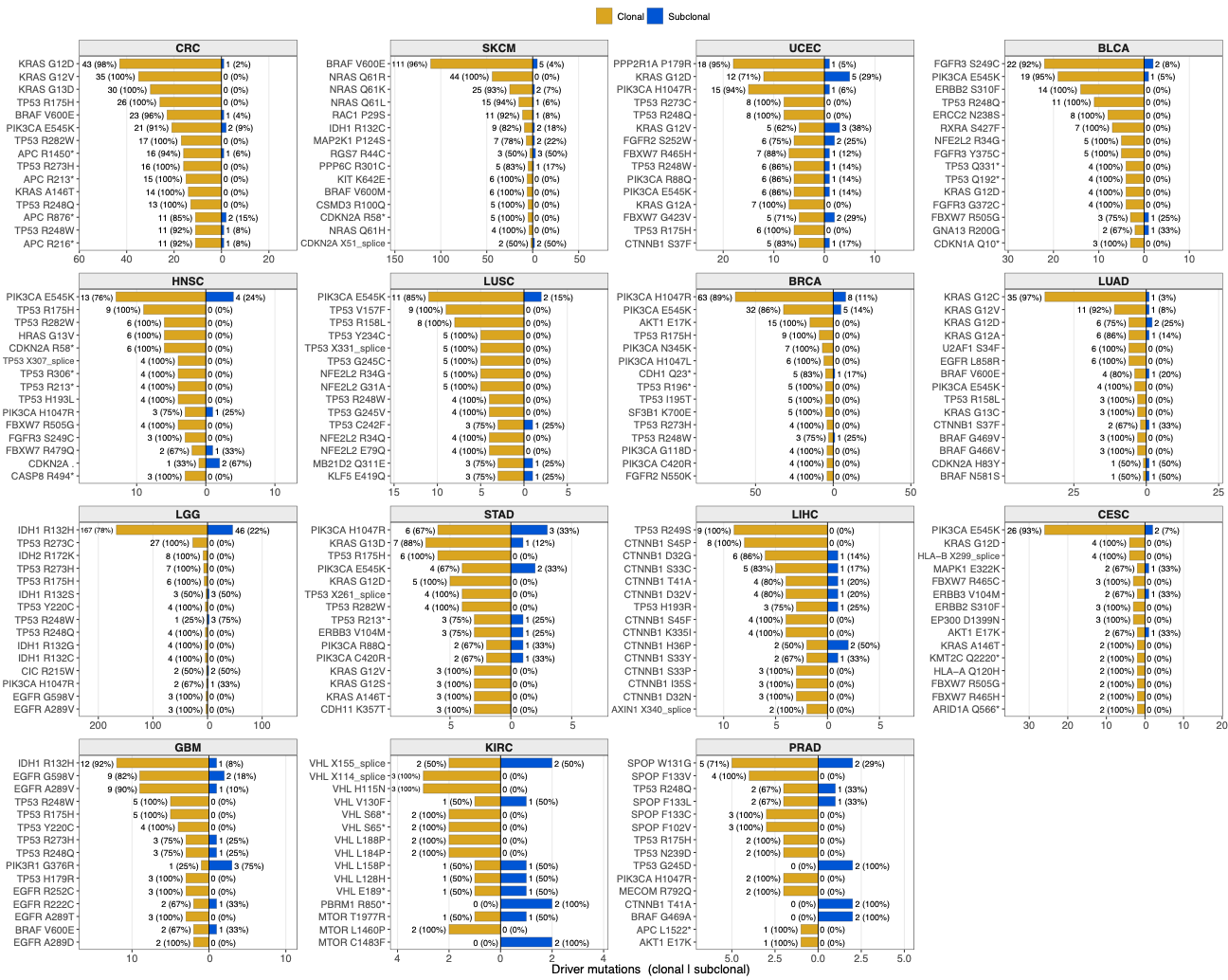

**Supplementary Figure S6: Cancer-type-specific clonality of recurrent driver hotspots.** For each of the 15 TCGA cancer types in the high-confidence cohort in which at least one recurrent locus carried two or more subclonal driver mutations, up to 15 of the most frequently observed hotspot loci are shown. Each row represents a gene and protein-change combination. Mirrored bars show the numbers of tumors in which the hotspot was classified as clonal (gold, left) or subclonal (blue, right) by CliPP. Labels report the tumor count and the corresponding percentage among tumors carrying that hotspot within the indicated cancer type. Hotspots are ordered by their total mutation count within each panel, and horizontal axes are scaled independently across cancer types.

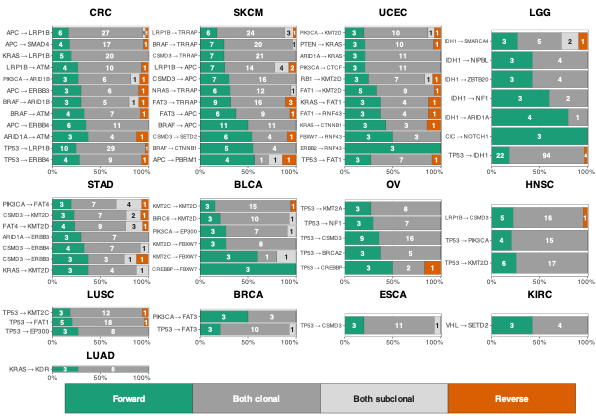

**Supplementary Figure S7. Cancer-type-specific relative timing of co-mutated driver pairs.** Co-mutated driver-gene pairs are summarized within each TCGA cancer types according to their joint clonal states. For a pair labeled A → B, Forward indicates that A was clonal and B was subclonal and is interpreted as A preceding B; Reverse indicates that A was subclonal and B was clonal and is interpreted as B preceding A. Both clonal and Both subclonal denote configurations for which relative order cannot be resolved. Eligible pairs were required to have at least 3 tumors with resolvable relative timing (Forward or Reverse), with these tumors accounting for at least 20% of all tumors carrying the pair. Among eligible pairs, the 12 most frequent pairs in each cancer type, ranked by the total number of co-mutated tumors, are displayed. Stacked bars show the proportions of co-mutated tumors classified as Forward (green), Both clonal (dark grey), Both subclonal (light grey), or Reverse (orange); values within segments indicate tumor counts.
