## Additional File 2 for "Scalable subclonal reconstruction of cancer cells in DNA sequencing data using a penalized likelihood model"

### **Additional file 2: Supplementary mathematical derivations and computational details**

### **1. Technical considerations for CliPP**

*Notations:*

| Parameter | Definition |
| --- | --- |
| $\boldsymbol{\rho}$ | Tumor purity, takes value between 0 and 1 |
| $\boldsymbol{\beta}$ | Cancer cell fraction (CCF), takes value between 0 and 1 |
| $\boldsymbol{\phi}$ | Cellular prevalence (CP), is equal to $\boldsymbol{\rho\beta}$, takes value between 0 and 1 |
| $\boldsymbol{r}_{\boldsymbol{i}}$ | Number of reads observed with variant alleles covering SNV $i$ |
| $\boldsymbol{n}_{\boldsymbol{i}}$ | Total number of reads covering SNV $i$ |
| $\boldsymbol{c}_{\boldsymbol{i}}^{\boldsymbol{T}}$ | Total copy number for tumor cell covering SNV $i$ |
| $\boldsymbol{c}_{\boldsymbol{i}}^{\boldsymbol{N}}$ | Total copy number for normal cell covering SNV $i$ |
| $\boldsymbol{m}_{\boldsymbol{i}}^{\boldsymbol{T}}$ | Copy number of the major allele covering SNV $i$ |
| $\boldsymbol{b}_{\boldsymbol{i}}^{\boldsymbol{V}}$ | SNV-specific copy number covering SNV $i$, also known as multiplicity |
| $\boldsymbol{\lambda}$ | Tuning parameter that controls the degree of the penalization |
| $\boldsymbol{S}$ | Total number of SNVs |
| $\boldsymbol{D}$ | Average read depth for the sample |
| $\boldsymbol{\theta}$ | Expected proportion of the variant allele |

#### **1.1 Computational structure using ADMM**

The alternating direction method of multipliers (ADMM) is a well-established algorithm that addresses constrained optimization challenges by decomposing them into simpler subproblems.

To solve the optimization problem in Eq.(8) in the manuscript, we introduce auxiliary variables $\eta_{ij}=\omega_{i}-\omega_{j}$ for $i<j$, thereby reformulating the problem as an equivalent constrained optimization problem:

|  | $S\left( \boldsymbol{\omega,\eta}; \lambda\right)=-\hat{\mathcal{l}}\left( \boldsymbol{\omega} \right)+\sum_{1\leq i<j\leq S} p_{\lambda}\left( \left\vert\eta_{ij} \right\vert\right)$, subject to $\omega_{i}-\omega_{j}-\eta_{ij}=0.$ | (S.1) |
| --- | --- | --- |

Using the augmented Lagrangian method, we further translate (S.1) into an unconstrained optimization problem and obtain the estimation of parameters by minimizing:

|  | $L\left( \boldsymbol{\omega,\eta,\tau}; \lambda\right)=S\left( \boldsymbol{\omega,\eta}; \lambda\right)+\frac{\alpha}{2}\sum_{i<j} \left( \omega_{i}-\omega_{j}-\eta_{ij} \right)^{2}-\sum_{i<j} \tau_{ij}\left( \omega_{i}-\omega_{j}-\eta_{ij} \right),$ | (S.2) |
| --- | --- | --- |

where the dual variables $\boldsymbol{\tau}=\left\{ \tau_{ij}, i<j \right\}$ are Lagrange multipliers, and $\alpha>0$ is the augmented Lagrangian penalty parameter. The implementation initializes $\alpha$ at 0.8 for each $\lambda$ and multiplies it by 1.02 after each ADMM iteration. During the update from iteration $k$ to $k+1$, $\alpha$ denotes the current value $\alpha_{k}$. The estimates of $\boldsymbol{\omega,\eta}$, and $\boldsymbol{\tau}$ are then computed iteratively by ADMM. The bottleneck in updating $\boldsymbol{\omega}$ at each ADMM iteration lies in the complex function $-\hat{\mathcal{l}}\left( \boldsymbol{\omega} \right)$. To address this challenge, we first use an estimated variance for the normal distribution and then employ a three-part piece-wise linear function to approximate each $g(\omega{}_{i})$, leading to a quadratic approximation to $-\hat{\mathcal{l}}\left( \boldsymbol{\omega} \right)$ (see **Sections 1.2 and 1.3 for the full derivation**). It is noteworthy that when $\gamma>\frac{1}{\alpha}+ 1$, the objective function $L\left( \boldsymbol{\omega, \eta, \tau}; \lambda\right)$ based on the SCAD penalty is convex with respect to $\eta_{ij}$ [1]. Furthermore, the minimizer of $L\left( \boldsymbol{\omega, \eta, \tau}; \lambda\right)$ with respect to $\eta_{ij}$ is unique and has a closed-form expression for given $\left( \boldsymbol{\omega, \tau} \right)$. Therefore, updates of $\boldsymbol{\omega,\eta}$, and $\boldsymbol{\tau}$ at one ADMM iteration proceed as follows. Given $\boldsymbol{\omega}^{\left( k \right)}$, $\boldsymbol{\tau}^{\left( k \right)}$ and $\boldsymbol{\eta}^{\left( k \right)}$ obtained from the $k$-th iteration, the update $\boldsymbol{\omega}^{\left( k+1 \right)}$ is given by:

|  | $\boldsymbol{\omega}^{\left( k+1 \right)}=\left( \boldsymbol{B}^{T}\boldsymbol{B}+\alpha\boldsymbol{\Delta}^{\boldsymbol{T}}\boldsymbol{\Delta} \right)^{-1}[\boldsymbol{\Delta}^{T}\left( \alpha\boldsymbol{\eta}^{\left( k \right)}+\boldsymbol{\tau}^{\left( k \right)} \right)-\boldsymbol{B}^{T}\boldsymbol{A}],$ (S.3) |
| --- | --- |

where $\boldsymbol{A}$ is a vector associated with $\boldsymbol{\omega}^{\left( k \right)}$, $\boldsymbol{B}$ is a matrix associated with $\boldsymbol{\omega}^{\left( k \right)}$ (see **Section 1.3 for the full derivation**), and $\boldsymbol{\Delta}$ is an $\left[ \frac{S\left( S-1 \right)}{2} \right]\times S$ matrix with rows $\left( e_{i}-e_{j} \right)^{T}$ for $1\leq i<j\leq S$, with $e_{i}$ defined as an $S\times1$ vector whose $i$-th element is 1 and others are 0. For given $(\boldsymbol{\omega}^{\left( k+1 \right)}, \boldsymbol{\tau}^{\left( k \right)})$, $\eta_{ij}$ is updated by solving:

$$\hat{\eta}_{ij}=\mathrm{argmin}_{\eta_{ij}}\frac{\alpha}{2}\left( \delta_{ij}-\eta_{ij} \right)^{2}+p_{\lambda}\left( \left| \eta_{ij} \right| \right),$$

where $\delta_{ij}= \omega_{i}^{(k+1)}- \omega_{j}^{(k+1)}- \alpha^{-1} \tau_{ij}^{\left( k \right)}$. This is a one-dimensional SCAD-penalized linear regression problem which has a closed-form solution. Thus, update of $\eta_{ij}$ at the $\left( k+1 \right)$-th iteration for SCAD penalty with $\gamma>1/\alpha+1$ is given by Ma and Huang [1]

|  | $\hat{\eta}_{ij}=\left\{ \begin{aligned} ST\left( \delta_{ij},\lambda/\alpha\right), \mathrm{if} \left\vert\delta_{ij} \right\vert\leq\lambda+\lambda/\alpha, \\ \frac{ST(\delta_{ij},\gamma\lambda/\left( (\gamma-1 \right)\alpha))}{1-1/(\left( \gamma-1 \right)\alpha)}, \mathrm{if} \lambda+\lambda/\alpha<\left\vert\delta_{ij} \right\vert\leq\gamma\lambda\\ \delta_{ij}, &if \left\vert\delta_{ij} \right\vert>\gamma\lambda\end{aligned}, \right.$ (S.4) |
| --- | --- |

where $ST\left( x,t \right)$ is the soft thresholding operator defined as $\mathrm{sign}\left( x \right)\left( \left| x \right|-t \right)$ if $\left| x \right|\geq t$ and 0 otherwise. At last, we update $\boldsymbol{\tau}$ by:

|  | $\boldsymbol{\tau}^{\left( k+1 \right)}=\boldsymbol{\tau}^{\left( k \right)}-\alpha\left( \boldsymbol{\Delta}\boldsymbol{\omega}^{\left( k+1 \right)}-\boldsymbol{\eta}^{\left( k+1 \right)} \right).$ (S.5) |
| --- | --- |

Consequently, the ADMM algorithm to solve the proposed model can be summarized as follows: (i) Initialize the estimates. Normally, a random initialization works well. We initialize $\omega_{i}^{\left( 0 \right)}=\log(\phi_{i}^{\left( 0 \right)}/[{1-\phi}_{i}^{(0)}])$, $\eta_{ij}^{\left( 0 \right)}=\omega_{i}^{\left( 0 \right)}-\omega_{j}^{\left( 0 \right)}$, and $\boldsymbol{\tau}^{\left( 0 \right)}=\boldsymbol{1}_{S(S-1)/2}$, where $1_{S(S-1)/2}$ is a vector of 1's with length $S(S-1)/2$. To obtain $\phi_{i}^{\left( 0 \right)}$, we first invert Eq. (1) in the main text at $\theta_{i}=r_{i}/n_{i},$with $c_{i}^{N}=2$. We divide the resulting CP estimates by the larger of 1 and their maximum, then clip them to $[\varepsilon,1-\varepsilon]$, where $\varepsilon=\left( 1+e^{5} \right)^{-1}$. (ii) At the $(k+1)$-th iteration, compute $\left( \boldsymbol{\omega}^{\left( k+1 \right)}, \boldsymbol{\eta}^{\left( k+1 \right)}, \boldsymbol{\tau}^{\left( k+1 \right)} \right)$ as described above. (iii) Terminate the algorithm if a stopping criterion (e.g., $\left\| \boldsymbol{\eta}^{\left( k+1 \right)}-\boldsymbol{\eta}^{\left( k \right)} \right\|_{\infty}<0.01$) is met. Otherwise, go back to step (ii) and repeat.

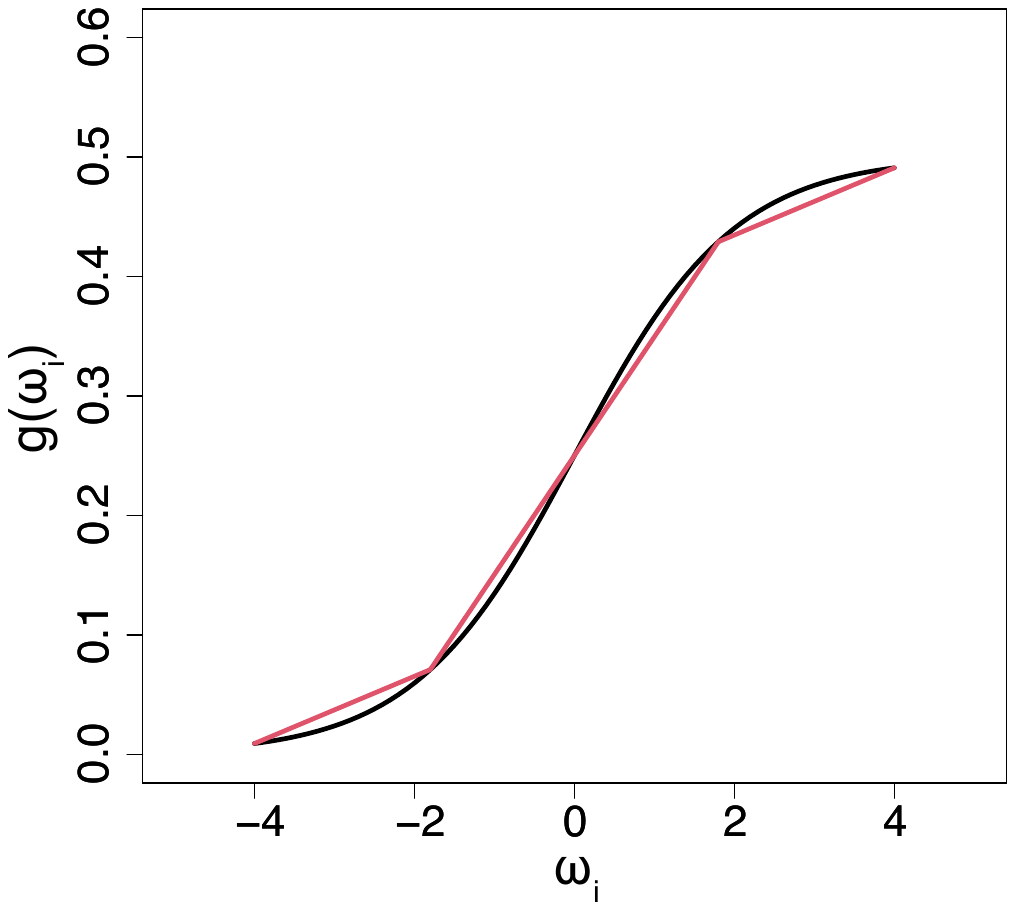

#### **1.2 Brief derivation of the linear approximation**

To address the challenge in updating $\boldsymbol{\omega}$, we adopt a three-part piece-wise linear function to approximate $g(\omega{}_{i})$ in the negative log-likelihood in Eq.(8) in the Method section. Shown on the right is an example of such an approximation. Specifically, we assume the linear approximation takes the form of $g(\omega_{i}) \approx u_{i} + v_{i}\omega_{i}.$ At iteration $k+1$, $u_{i}$ and $v_{i}$ are the intercept and slope, respectively, of the linear segment containing $\omega_{i}^{k}$, and are held fixed during the update of $\omega$. Recall in the manuscript, we already have:

$$\theta_{i}=g\left( \omega_{i} \right)=\frac{b_{i}^{V}e^{\omega_{i}}}{(1+e^{\omega_{i}})A}$$

where $A=2-2\rho+c_{i}^{T}\rho$ is a constant. Let’s set

$$f\left( x \right)=\frac{b_{i}^{V}e^{x}}{\left( 1+e^{x} \right)A}$$

$$g\left( x \right)=\left\{ \begin{aligned} a_{1}x+b_{1}, \mathrm{if} x\leq x_{1} \\ a_{2}x+b_{2}, \mathrm{if} x\geq x_{2} \\ a_{3}x+b_{3}, {\mathrm{if} x}_{1}<x<x_{2} \end{aligned} \right.$$

where $g(x)$ is used to approximate$f\left( x \right).$ We define the approximation error $D$as follows:

$$D=\sup_{x\in G} |f\left( x \right)-g(x)|$$

where $G\subset[-M,M]$ is a finite evaluation grid containing both endpoints, with ($M>0$). Then our goal is to find a pair of separation points $\left( x_{1},x_{2} \right)$ for constructing $g(x)$ that achieves the smallest $D$. To that end, we employ a grid search strategy and limit the range of $x$ to $[-M, M]$ to save computational cost. In practice, we set $M=4$ which can roughly recover the full dynamic range of $f\left( x \right)$. The grid spans from $-M$ to $M$ with a step size of 0.1, and we record the approximation error $D$ at each pair of separation points $\left( x_{1},x_{2} \right)$. Finally, the approximation function is designed using the pair of $\left( x_{1},x_{2} \right)$ associated with the smallest $D$.

#### **1.3 Computational details**

To estimate $\boldsymbol{\omega, \eta, \tau}$, we designed an ADMM-based algorithm for minimizing the objective function in Eq.(S.2). Given $(\boldsymbol{\eta, \tau})$, an update of $\boldsymbol{\omega}$ can be derived by setting $\partial L(\boldsymbol{\omega, \eta, \tau} ; \lambda)/\partial\boldsymbol{\omega}=0$. However, it is difficult to directly compute the derivative. Here, we utilize two strategies. First, instead of using the theoretic variance, we use an estimated variance based on the estimated $\boldsymbol{\omega}$ from the previous step. Second, we approximate $g(\omega{}_{i})$ by a linear function as discussed above in **Section 1.2**. That is, at step $k+1$, we rewrite the objective function as follows:

$L\left( \boldsymbol{\omega,\eta,\tau};\lambda,\boldsymbol{\omega}^{\left( k \right)} \right)=S\left( \boldsymbol{\omega,\eta};\lambda, \boldsymbol{\omega}^{\left( k \right)} \right)+\frac{\alpha}{2}\sum_{i<j} \left( \omega_{i}-\omega_{j}-\eta_{ij} \right)^{2}-\sum_{i<j} \tau_{ij}\left( \omega_{i}-\omega_{j}-\eta_{ij} \right)$

|  | $=\sum_{i=1}^{S} \{-log\frac{1}{\sqrt{2\pi n_{i}g\left( \omega_{i} \right)\left( 1-g\left( \omega_{i} \right) \right)}}+\frac{\left( n_{i}g\left( \omega_{i} \right)-r_{i} \right)^{2}}{2n_{i}g\left( \omega_{i} \right)\left( 1-g\left( \omega_{i} \right) \right)}\}$+$\sum_{1\leq i<j\leq S} p_{\lambda}\left( \left\vert\eta_{ij} \right\vert\right)$  $+\frac{\alpha}{2}\sum_{i<j} \left( \omega_{i}-\omega_{j}-\eta_{ij} \right)^{2}-\sum_{i<j} \tau_{ij}\left( \omega_{i}-\omega_{j}-\eta_{ij} \right)$ |
| --- | --- |
|  | $\approx\frac{1}{2}\sum_{i=1}^{S} \frac{n_{i}\left( g\left( \omega_{i} \right)-\frac{r_{i}}{n_{i}} \right)^{2}}{g\left( \omega_{i}^{\left( k \right)} \right)\left( 1-g\left( \omega_{i}^{\left( k \right)} \right) \right)}+\sum_{1\leq i<j\leq S} p_{\lambda}\left( \left\vert\eta_{ij} \right\vert\right)+\frac{\alpha}{2}\sum_{i<j} \left( \omega_{i}-\omega_{j}-\eta_{ij} \right)^{2}-\sum_{i<j} \tau_{ij}\left( \omega_{i}-\omega_{j}-\eta_{ij} \right)+C_{0}$ |
|  | $=\frac{1}{2}\sum_{i=1}^{S} \frac{n_{i}\left( u_{i}+v_{i}\omega_{i}-\frac{r_{i}}{n_{i}} \right)^{2}}{g\left( \omega_{i}^{\left( k \right)} \right)\left( 1-g\left( \omega_{i}^{\left( k \right)} \right) \right)}+\sum_{1\leq i<j\leq S} p_{\lambda}\left( \left\vert\eta_{ij} \right\vert\right)+\frac{\alpha}{2}\sum_{i<j} \left( \omega_{i}-\omega_{j}-\eta_{ij} \right)^{2}-\sum_{i<j} \tau_{ij}\left( \omega_{i}-\omega_{j}-\eta_{ij} \right)+C_{0}$ |
|  | $=\frac{1}{2}\sum_{i=1}^{S} \left( A_{i}+B_{i}\omega_{i} \right)^{2}+\frac{\alpha}{2}\sum_{i<j} \left( \left( e_{i}-e_{j} \right)^{T}\boldsymbol{\omega}-\eta_{ij}-\alpha^{-1}\boldsymbol{\tau}_{\boldsymbol{ij}} \right)^{2}+C$ |
|  | $=\frac{1}{2}\left\vert\left\vert\boldsymbol{A}+\boldsymbol{B\omega} \right\vert\right\vert^{2}+\frac{\alpha_{k}}{2}\left\vert\left\vert\Delta\boldsymbol{\omega}-\boldsymbol{\eta}-\alpha_{k}^{-1}\boldsymbol{\tau} \right\vert\right\vert^{2}+C$ |

where $A_{i}=\frac{\sqrt{n_{i}}(u_{i}-r_{i}/n_{i})}{\sqrt{g(\omega_{i}^{\left( k \right)})(1-g(\omega_{i}^{(k)}))}}$, and $B_{i}=\frac{\sqrt{n_{i}}v_{i}}{\sqrt{g(\omega_{i}^{\left( k \right)})(1-g(\omega_{i}^{(k)}))}}$,$C$ is a function independent of $\boldsymbol{\omega}$, $e_{i}$ is $S\times1$ vector whose $i^{th}$ element is 1 and others are 0, and $\Delta=\left\{ \left( e_{i}-e_{j} \right), i<j \right\}^{T}$, A is a vector given by $\boldsymbol{A}=\left\{ A_{1}, ..., A_{S} \right\}^{T}$, and $\boldsymbol{B}$ is a matrix defined as

$$\boldsymbol{B}=\left[ \begin{matrix} B_{1} & 0 & \cdots& 0 \\ 0 & B_{2} & \cdots& 0 \\ \vdots& \vdots& \ddots& \vdots\\ 0 & \cdots& 0 & B_{S} \end{matrix} \right]$$

Therefore,

$$\boldsymbol{\omega}^{\left( k+1 \right)}=\left( \boldsymbol{B}^{T}\boldsymbol{B}+\alpha\boldsymbol{\Delta}^{\boldsymbol{T}}\boldsymbol{\Delta} \right)^{-1}\left[ \boldsymbol{\Delta}^{T}\left( \alpha\boldsymbol{\eta}^{\left( k \right)}+\boldsymbol{\tau}^{\left( k \right)} \right)-\boldsymbol{B}^{T}\boldsymbol{A} \right].$$

The direct inversion of matrix $\boldsymbol{B}^{T}\boldsymbol{B} +\alpha\Delta^{T}\Delta$ may not be feasible when $S$ is large, but fortunately we do not need to perform an $S\times S$ matrix inversion, as Miller [2] suggested. Notice that $\boldsymbol{\Delta}^{\boldsymbol{T}}\boldsymbol{\Delta} = S\boldsymbol{I}_{S}-\boldsymbol{1}_{S}\boldsymbol{1}_{\boldsymbol{S}}^{T}$, thus,

$\boldsymbol{B}^{T}\boldsymbol{B}+\alpha\Delta^{T}\Delta=\left( \boldsymbol{B}^{T}\boldsymbol{B+}S\alpha\boldsymbol{I}_{S} \right)-\alpha\boldsymbol{1}_{S}\boldsymbol{1}_{S}^{\boldsymbol{T}}$.

Now let $\boldsymbol{M= B}^{T}\boldsymbol{B+}S\alpha\boldsymbol{I}_{S},$ and a rank-one correction $\boldsymbol{R}=-\alpha\boldsymbol{1}_{S}\boldsymbol{1}_{S}^{\boldsymbol{T}}$**.** $\boldsymbol{M}$ is easy to invert since it is a diagonal matrix with non-zero diagonal elements, and $rank(\boldsymbol{R})=1$. Then we have:

$$\left( \boldsymbol{B}^{T}\boldsymbol{B}+\alpha\boldsymbol{\Delta}^{\boldsymbol{T}}\boldsymbol{\Delta} \right)^{-1}=\left( \boldsymbol{M}+\boldsymbol{R} \right)^{-1}=\boldsymbol{M}^{-1}-\frac{1}{1+g}\boldsymbol{M}^{-1}\boldsymbol{R}\boldsymbol{M}^{-1}$$

where $g = trace\left( \boldsymbol{R}\boldsymbol{M}^{-1} \right).$ Here we have avoided computing the inverse of an $S\times S$ matrix, instead, we only need to compute a series of matrix multiplications. We utilize graphics processing unit (GPU) computing for further acceleration. In the right side of equation above, matrix operations $\boldsymbol{M}^{-1}$ and $\boldsymbol{M}^{-1}\boldsymbol{R}\boldsymbol{M}^{-1}$ are GPU-friendly. Note that $\boldsymbol{M}$ is a diagonal matrix which leads to $\boldsymbol{M}^{-1}$ is diagonal matrix and able to compute independently for each SNV. In iteration $k+1$, the update $\boldsymbol{\omega}^{\left( k+1 \right)}$ can be decomposed and distributed across thousands of GPU threads. Therefore, the GPU accelerates the algorithm by parallelizing the dominant $O(S^{2})$ pairwise ADMM operations. The updates of $\boldsymbol{\eta}$ and $\boldsymbol{\tau}$ are achieved by Eq.(S.4) and Eq.(S.5), respectively, as discussed in **Section 1.1**.

**2.** **Advances in CliPP 1.4.0/1.4.1**

CliPP has been substantially revised since the early development version used in the Dentro et al. 2021 PCAWG consensus [3]. Details of the current implementation are described above, with major changes highlighted here. The version 1.4.0 and 1.4.1 releases feature corrected multiplicity computation and a reformulated penalized likelihood, along with the current improved regularization hyperparameter selection. Both the preprocessing and postprocessing steps were rewritten, fixing bugs that had produced incorrect cluster assignments for some mutations. In CliPP 1.4.1 the pairwise computations, which dominate runtime and scale quadratically with the number of mutations, were ported to CUDA [4], giving the option to use accelerated GPU computation when there are large numbers of SNVS, which results in an approximately 10x speed improvement on WGS samples. Further improvements have also been made to facilitate user implementation, with install now available from Github or from a docker image, and the entire pipeline being runnable with a single command.
